# The Developmental Transcriptome of *Ae. albopictus,* a Major Worldwide Human Disease Vector

**DOI:** 10.1101/753962

**Authors:** Stephanie Gamez, Igor Antoshechkin, Stelia C. Mendez-Sanchez, Omar S. Akbari

**Affiliations:** Section of Cell and Developmental Biology, University of California, San Diego, La Jolla, California, United States of America; Tata Institute for Genetics and Society-UCSD, La Jolla, California, USA; Division of Biology and Biological Engineering, California Institute of Technology, Pasadena, California, 91125, USA; Group for Research in Biochemistry and Microbiology (Grupo de Investigación en Bioquímica Y Microbiología-GIBIM), School of Chemistry, Universidad Industrial de Santander, Bucaramanga, Colombia

**Keywords:** *Aedes albopictus*, transcriptome, development, Zika, dengue, *Aedes aegypti*, RNA-seq

## Abstract

*Aedes albopictus* mosquitoes are important vectors for a number of human pathogens including the Zika, dengue, and chikungunya viruses. Capable of displacing *Aedes aegypti* populations, it adapts to cooler environments which increases its geographical range and transmission potential. There are limited control strategies for *Aedes albopictus* mosquitoes which is likely attributed to the lack of comprehensive biological studies on this emerging vector. To fill this void, here using RNAseq we characterized *Aedes albopictus* mRNA expression profiles at 47 distinct time points throughout development providing the first high-resolution comprehensive view of the developmental transcriptome of this worldwide human disease vector. This enabled us to identify several patterns of shared gene expression among tissues as well as sex-specific expression patterns. Moreover, to illuminate the similarities and differences between *Aedes aegypti*, a related human disease vector, we performed a comparative analysis using the two developmental transcriptomes. We identify life stages were the two species exhibited significant differential expression among orthologs. These findings provide insights into the similarities and differences between *Aedes albopictus* and *Aedes aegypti* mosquito biology. In summary, the results generated from this study should form the basis for future investigations on the biology of *Aedes albopictus* mosquitoes and provide a goldmine resource for the development of transgene-based vector control strategies.

## Introduction

*Aedes albopictus* (*Ae. albopictus*), the Asian tiger mosquito, is a medically important invasive species whose habitat range has significantly increased over the past 20 years (Benedict *et al.* 2007). Originally from East Asia and the islands of the Pacific and Indian Ocean, this species is now found in all continents except for Antarctica (Bonizzoni *et al.* 2013). The rapid expansion of *Ae. albopictus* has been attributed to its ecological plasticity, strong competitive aptitude, feeding behavior, vector competence and its ability to enter diapause to escape unfavorable seasonal conditions, and lack of effective control strategies (Reynolds *et al.* 2012). An increase in habitat range imposes a greater risk of transmitting several mosquito-borne pathogens such as Zika, chikungunya, and dengue virus (Rezza 2012; Shragai *et al.* 2017). Even though this species is considered a less efficient dengue vector than *Aedes aegypti* (*Ae. aegypti*), the primary vector of dengue virus and a closely related mosquito species, it is responsible for several outbreaks of dengue and chikungunya virus (Vazeille *et al.* 2007; Ratsitorahina *et al.* 2008; Pagès *et al.* 2009).

Today both *Ae. albopictus* and *Ae. aegypti* are found in most Asian cities and in large parts of the Americas (Lambrechts *et al.* 2011). Both *Ae. albopictus* and *Ae. aegypti* feed on humans in daylight hours and rest indoors (Harrington *et al.* 2001; Valerio *et al.* 2010; Dzul-Manzanilla *et al.* 2016) and have similar larval niches, but their distributions depend on local environmental conditions (Kraemer *et al.* 2019). Interestingly, there are differences in dominance of mosquito vectors along urban-rural gradients. *Ae. albopictus* is often found in urban and rural environments whereas *Ae. aegypti* tends to be an urban vector utilizing artificial containers (Tsuda *et al.* 2006; Kraemer *et al.* 2015). When both populations of mosquitoes are present in the same ecological niche *Ae. albopictus* mosquitoes tend to outcompete *Ae. aegypti* (O’meara *et al.* 1995; Beilhe *et al.* 2012). This is hypothesized by *Ae. albopictus* being a superior larval competitor (Bagny Beilhe *et al.* 2013). In addition, *Ae. albopictus* is found to be ecologically plastic where it can survive in cooler environments than *Ae. aegypti*, thus facilitating its spread to unconventional environments (Kraemer *et al.* 2019). To help combat this emerging mosquito, we need a better understanding of its biology to enable the innovation of effective control strategies.

Previously, a comprehensive developmental transcriptome study of *Ae. aegypti* analyzed and provided insight into the complexity of the basic biology of these mosquitoes (Akbari *et al.* 2013a; Matthews *et al.* 2018). This enormous dataset has provided the community with a foundation of data enabling the functional characterization of novel germline promoters (Akbari *et al.* 2014b) which have subsequently been used to develop highly potent Cas9 endonuclease expressing strains (Li *et al.*; Akbari *et al.* 2014b) in addition to gene drives (Li *et al.* 2019). While diapause has been studied extensively for *Ae. albopictus* because of its importance to its survival in different environments (Urbanski *et al.* 2010a, 2010b; Reynolds *et al.* 2012; Diniz *et al.* 2017), and the genome has been sequenced (Chen *et al.* 2015), there is currently no developmental transcriptome available for *Ae. albopictus*. Therefore to fill this void, here we provide a comprehensive analysis of the *Ae. albopictus* transcriptome throughout development sequenced at unprecedented depth which will provide the community with a goldmine of data and also a unique opportunity to perform comparative analysis and may even enable the discovery of novel genes and regulatory elements which may prove useful for innovating genetic control strategies.

## Results

### *Ae. albopictus* developmental transcriptome timepoints

To establish a comprehensive global view of gene expression dynamics throughout *Ae. albopictus* development, we performed Illumina RNA sequencing (RNA-Seq) on 47 unique samples representing 34 distinct stages of development. These time points incorporated 31 whole animal and 16 tissue/carcass samples. For example, for embryogenesis 19 samples were collected; the first three time points, 0-1 hr, 0-4 hr, and 4-8 hr embryos, capture the maternal-zygotic transition, whereas 16 additional embryo samples were collected at 4 hr intervals until 72 hr to capture the rest of embryogenesis. Samples from four larval stages and sex-separated early and late male and female pupae were collected to capture the aquatic life cycle. Additionally, whole dissected ovaries and carcasses (whole female bodies lacking ovaries) from non-blood-fed females (NBF) and from females at 12 hr, 24 hr, 36 hr, 48 hr, 60 hr, and 72 hr post-blood-meal (PBM) were collected to examine the pre-vitellogenic “resting stage” through the completion of oogenesis. Adult male testes and carcasses (lacking testes) were collected at four days post eclosion to investigate male-specific germline and somatic gene expression. A complete list of the samples collected is provided in Supplemental Table S1. To achieve single nucleotide resolution we extracted total RNA from each sample and sequenced entire transcriptomes using the Illumina HiSeq2500, and generated 1.56 billion 50nt reads corresponding to total sequence output of 78.19 GB with close to 95% of the reads aligning to the most contiguous and complete *Ae. albopictus* assembly available (assembly: canu_80X_arrow2.2, strain: C6/36, VectorBase) (Supplemental Fig. S1A; Supplemental Tables S2, S3, S4).

### Global transcriptome dynamics

To capture the global dynamics of gene expression, we quantified the gene expression profiles across all developmental timepoints (Fig. 1; Supplemental Table S4). In general, the number of expressed genes (FPKM>1) gradually increases through embryogenesis, reaching a peak at 68-72 hr (Supplemental Fig. S1B). This likely reflects the embryo developing and preparing for the next major developmental stage, the larval stage. In this stage, the number of genes expressed are lower in 1^st^ larval instar, but then increases in the subsequent 2^nd^ to 4^th^ instars. After the larval stage, the animal undergoes metamorphosis into pupae where sexual dimorphism is prevalent. During the early pupal stages, the number of genes increases suggesting transcripts involved in hormone production and initiation of adult formation are being expressed (Margam *et al.* 2006). In the adult stages, the difference between the male and female germline is obvious, males express the highest number of genes. When females take a blood meal for egg production, the number of genes expressed in the ovary do not seem to vary significantly, however when looking at their corresponding carcasses, varying levels across PBM females is observed suggesting dynamic gene expression in the soma (Supplemental Fig. S1B). Interestingly, the tissue with the highest number of genes expressed (FPKM>1) corresponds to the male testes, while those with the lowest corresponds to 24 hr PBM female carcasses. Analysis of pairwise correlations revealed that almost every developmental stage is most highly correlated with its adjacent stage, particularly evident during embryogenesis (Figure 1A). Notable exceptions to this trend occur in 24-36 hr PBM female carcass and 36-48 hr PBM ovaries, suggesting that these represent important points where physiological transitions occur in blood-fed females. Diapause samples (0 through 4 weeks) and NBF and 12 hr PBM ovaries are highly correlated with the mid-stages of embryogenesis suggesting similar genes are expressed in these samples. In the 0-1 hr embryo time point, we see similar gene expression with the 60- and 72 hr PBM ovaries likely reflective of maternally deposited transcripts. Samples with unique gene expression include the male germline, 24 hr PBM female carcass, 24 hr and 36 hr PBM ovaries, and late pupae (Fig. 1A).

**Figure 1.**
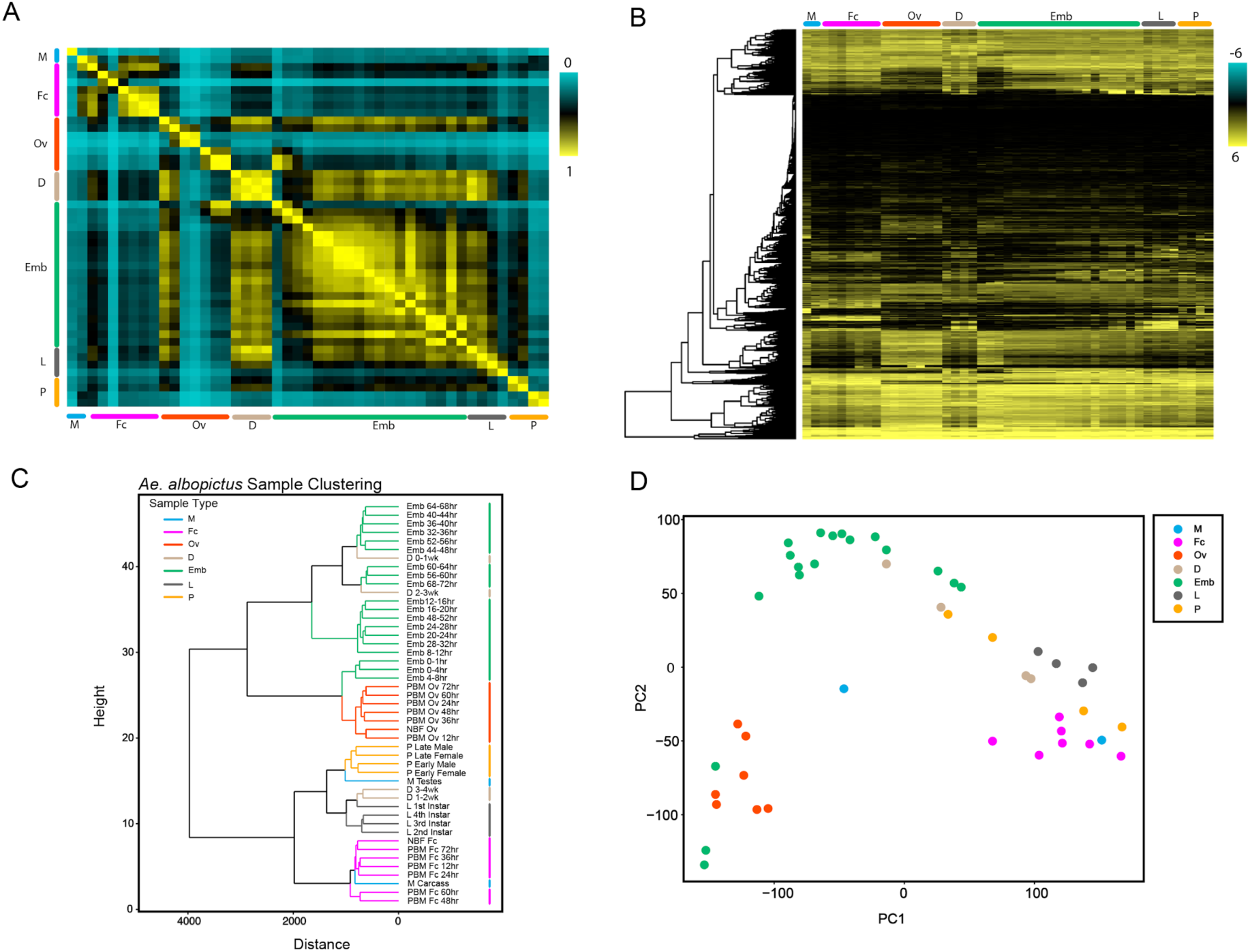
Global dynamics of gene expression. (A) Correlation matrix of all RNA seq timepoints for all known *Ae. albopictus* genes. (B) Hierarchical clustering heat map of albopictus genes across all developmental stages. FPKM values were log2(x+1) transformed and were scaled to plot the z-scores. (C) Dendrogram of *Ae. albopictus* samples clustering similar life stages closer together. Plot depicts the close relationship between all developmental samples. (D) PCA clustering of *Ae. albopictus* samples depicts clustering of life stages who show close similarity. PCA plot is in agreement with clustering dendrogram. For A-D, the major developmental groups are indicated by color bars and are organized as follows: M (blue, male testes, male carcass), Fc (pink, NBF carcass, and multiple timepoints PBM: 12hr, 24hr, 36hr, 48hr, 60hr, and 72hr), Ov (orange, NBF ovaries, and multiple ovarian timepoints PBM: 12hr, 24hr, 36hr, 48hr, 60hr, and 72hr), D (tan, diapause at multiple timepoints: 0-1wk, 1-2wk, 2-3wk, and 3-4wk), Emb (embryo at multiple timepoints: 0-1hr, 0-2hr, 2-4hr, 4-8hr, 8-12hr, 12-16hr, 16-20hr, 20hr-24hr, 24-28hr, 28-32hr, 32-36hr, 36-40hr, 40-44hr, 44-48hr, 48-52hr, 52-56hr, 56-60hr, 60-64hr, 64-68hr, and 68-72hr embryos), L (gray, larvae 1st, 2nd, 3rd, and 4th instar larvae stages), and P (yellow, pupae, early male and female, and late male and female pupae stages).

To further visualize the various patterns of gene expression and the relationships between the samples, hierarchical clustering and PCA analyses was performed (Fig. 1B, C, D). Based on these analysis, embryo, PBM ovaries, pupae, larvae, and PBM female carcass samples tend to cluster closer together which is expected as their gene expression profiles are similar as these are developmentally related samples. Interestingly however, in Figure 1D, the male testes sample clusters away from all other samples, reflecting a distinguishing difference between this sample as compared to other samples sequenced. To further visualize the patterns of co-regulated gene expression we used a soft clustering algorithm and identified 20 distinct patterns that included 543 to 2760 genes (Fig. 2A). Each cluster in Figure 2A contains a set of *Ae. albopictus* genes that have an assigned membership value to indicate the degree of similarity to genes in that cluster (Supplemental Table S5). The majority of these clustering patterns correspond to the developmental stages and transitions of the mosquito. For example, clusters 1 through 7 include genes that are associated with embryogenesis (Fig. 2A). To further investigate the functional associations of the genes in each cluster, we performed gene ontology (GO) analysis. Genes in clusters 1 through 7 are highly enriched in genes involving nucleic acid binding (e.g. LOC109417994, LOC109422308 and LOC109424406), organic cyclic compound binding (e.g. LOC109428278, LOC109432015 and LOC109431126), transcription factor activity for molecular function (e.g. LOC109397112, LOC109401789 and LOC109409855) macromolecule catabolic, metabolic, and biosynthetic processes (e.g. LOC109410609, LOC109410731 and LOC109429162), regulation of gene expression (e.g. LOC109399731, LOC109414325 and LOC109416300), signal transduction (e.g. LOC109413732), DNA-templated transcription, and regulation of nitrogen compound metabolic process (e.g. LOC109415851) for biological processes (Supplemental Table S5, S6). These processes correspond to the necessities of the developing embryo as it transitions between stages with rapidly changing demands. Energy is supplied to the embryo through the breakdown of biomolecules. A list of early zygotic and maternal genes are provided in Supplementary Tables S7 and S8, respectively, to depict the genetic profile of early embryogenesis. After embryogenesis, the deposited egg can enter diapause, a dormant state that allows the mosquito embryo to survive unfavorable conditions (Armbruster 2016). Cluster 8 includes gene expression required for diapause in dormant embryos. Here, several processes such as translation, protein metabolic, organonitrogen compound metabolic and biosynthetic processes are enriched as well as terms associated with lipid transporter activity, lipid-A-disaccharide synthase activity, ribonucleoside binding and structural molecular activity (Supplemental Table S6). Genes represented in this cluster include several lipases (to name a few: LOC109417138, LOC109401099, LOC109430899, and LOC109430905), several fatty acid hydroxylases (to name a few: LOC109400137, LOC109432075, and LOC109397180), and some proteases (LOC109406257, LOC109411917, and LOC109402104) (Supplemental Table S5). This is likely due to the expression of genes that correspond to specific metabolic events associated with diapause to enable cold tolerance (Diniz *et al.* 2017). When an embryo hatches under favorable conditions it then enters the aquatic larval stage. Clusters 9 and 10 correspond to these stages and have genes enriched for serine-type peptidase activity, chitin binding, metallopeptidase activity, oxidoreductase activity, and ATPase activity under the molecular function category. Biological processes taking place include proteolysis, amino sugar metabolic, chitin metabolic, glucosamine-containing compound metabolic, and amino sugar metabolic processes (Supplemental Table S6). The metabolic processes are likely involved in preparing the larva to acquire the energy reserves that will be used for egg development (Telang *et al.* 2006). Following the larval stage, the mosquito then enters the pupal stage, the final aquatic stage in the mosquito’s life cycle. Clusters 11, 12 and 15 include genes enriched for structural constituent of cuticle, oxidoreductase, peptidase, and serine-type peptidase activity, which are likely involved in immunity and the hydrolysis of nutrients (Saboia-Vahia *et al.* 2013). Steroid biosynthetic and metabolic processes are also prevalent which suggests hormones like ecdysteroids, which are crucial for metamorphosis, are preparing the pupa to molt into an adult mosquito (Margam *et al.* 2006).

**Figure 2.**
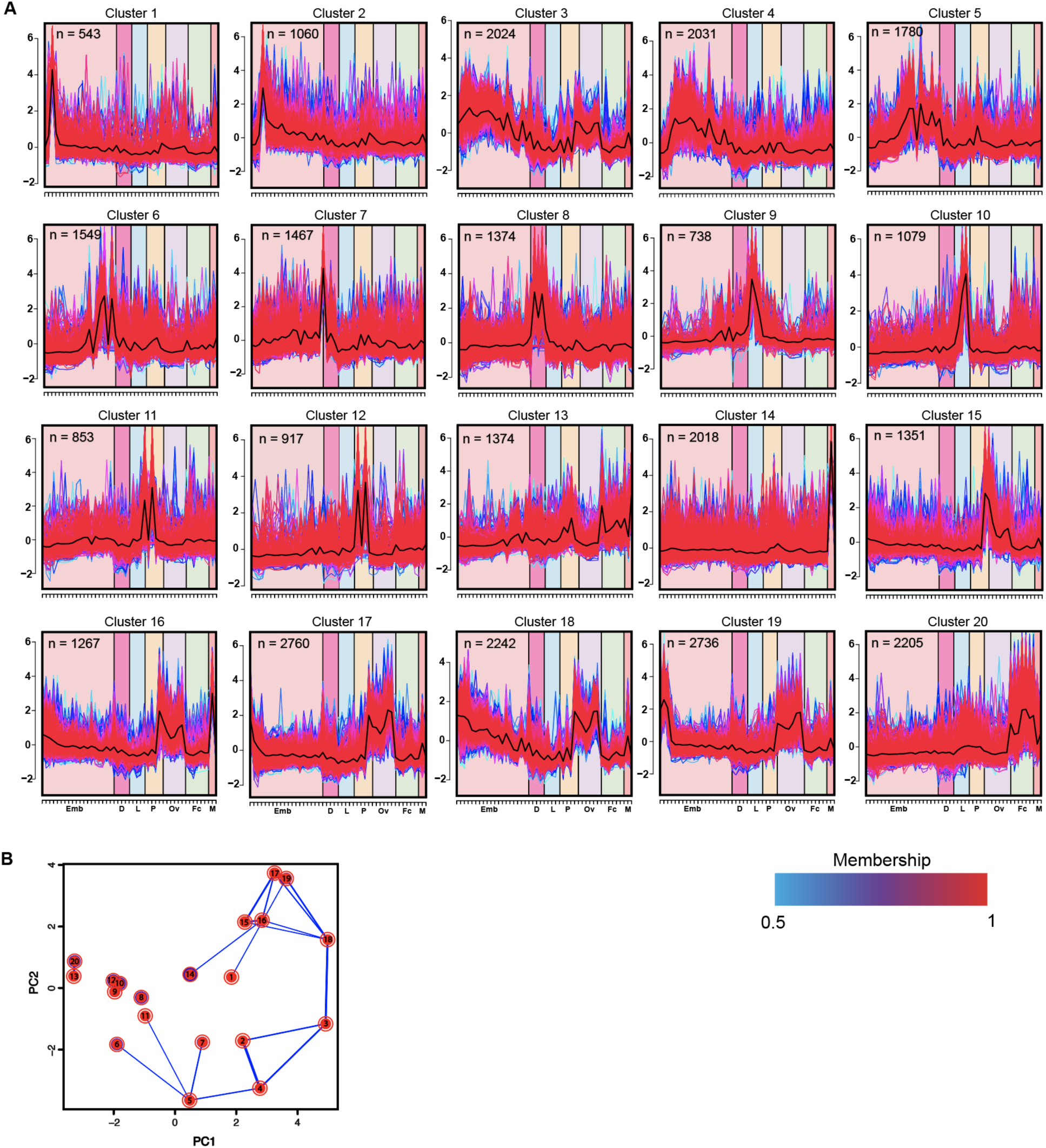
Soft clustering and principal component analysis on *Ae. albopictus* genes. (A) Twenty albopictus gene expression profile clusters were identified through soft clustering. Each gene is assigned a line color corresponding to its membership value, with red (1) indicating high association. The developmental groups are indicated by symbols on the x-axis. (B) Principal component analysis shows relationships between the 20 clusters, with thickness of the blue lines between any two clusters reflecting the fraction of genes that are shared. N, the number of genes in each cluster.

Following the pupal stage, the aquatic life cycle ends with the animal undergoing metamorphosis into an adult. In males, the carcass and gonads have different terms enriched (clusters 13 for carcass and 14 and 16 for male testes). The male somatic tissues (carcass) include terms enriched for neurotransmitter receptor activity, and several ATP biosynthetic process (Supplemental Table S6). Terms enriched in the male germline include microtubule motor activity, microtubule based processes, spermatid development, olfactory receptor activity, neurotransmitter receptor activity, and sensory perception of chemical stimulus. Like *Anopheles gambiae* and *Ae. aegypti* mosquitoes, *Ae. albopictus* use chemosensation to activate their spermatozoa by modulating sperm activation and perhaps the orientation of spermatozoa (Pitts *et al.* 2014). Clusters 17 through 19 correspond to blood-fed female ovaries and include genes enriched for cellular response to stimulus and Ras protein signal transduction which are important means of communication during the processes of oocyte and eggshell patterning (Dana *et al.* 2005). In addition, several metabolism processes that are crucial for the breakdown of organic molecules such as deoxyribonucleotides are crucial to support cell division (Telang *et al.* 2013). Finally, cluster 20 corresponds to the genes that are expressed in PBM female carcasses. Here, genes enriched for heme binding, lipid metabolic processes, sensory perceptions are all related to the processes required to support the future developing zygote in the ovaries. Other relevant genes enriched are those with serine-type endopeptidase activities, which is likely important for breaking down blood proteins (Bian *et al.* 2008).

### Sex specific gene expression overview

Genes expressed in the germline of males and females are believed to play an important role in evolution, contributing to reproductive fitness, isolation, and speciation (Whittle and Extavour 2017). Thus it is an important step to study the role of sex-biased gonad genes in evolution. To gain insight into the sex-specific differences in *Ae. albopictus*, we compared the transcriptomes of male and female samples (male testes and carcass, female non-blood-fed (NBF) and PBM ovaries and carcasses, and pupal samples) (Supplemental Fig. S2). When comparing NBF ovaries to male testes, we observed sex biased gene expression levels (out of 34,824 genes 550 and 67 genes were upregulated between testes and NBF ovary samples, respectively). Female-specific genes were determined by comparing genes that were differentially expressed between the male germline tissues and female germline tissues at several different PBM time points (12-, 24-, 36-, 48-, 60-, and 72hr). The same strategy was performed with the carcass samples to look at somatic-specific expression.

### Female-specific genes in NBF and PBM states

In our analysis, we found a total of 704 genes that were upregulated in the germline and somatic tissues of female mosquitoes and are listed in Supplemental Table S9. In female ovaries (NBF and PBM), several vitelline membrane proteins, chorion peroxidases, and uncharacterized genes are over expressed (Supplemental Fig. S3; Supplemental Table S9). Only six genes, four of which are uncharacterized and the other two correspond to an aminopeptidase M1-like gene and a trypsin-4-like gene, are specifically over-expressed in the NBF ovaries (Supplemental Table S9). Out of all these genes, only one (aminopeptidase M1-like) is found to be orthologous in *Ae. aegypti* while the other five seem to be specific to *Ae. albopictus*. This may be due to deficiencies in the annotation of the *Ae. albopictus* genome or may represent loci that are specific to this species. Additional work needs to be done to uncover the identity and function of these unknown loci that are highly expressed in the female ovaries. In NBF carcass samples, genes highly expressed include trypsins, 30kDa salivary gland allergens, and several uncharacterized genes (Supplemental Table S9).

**Figure 3.**
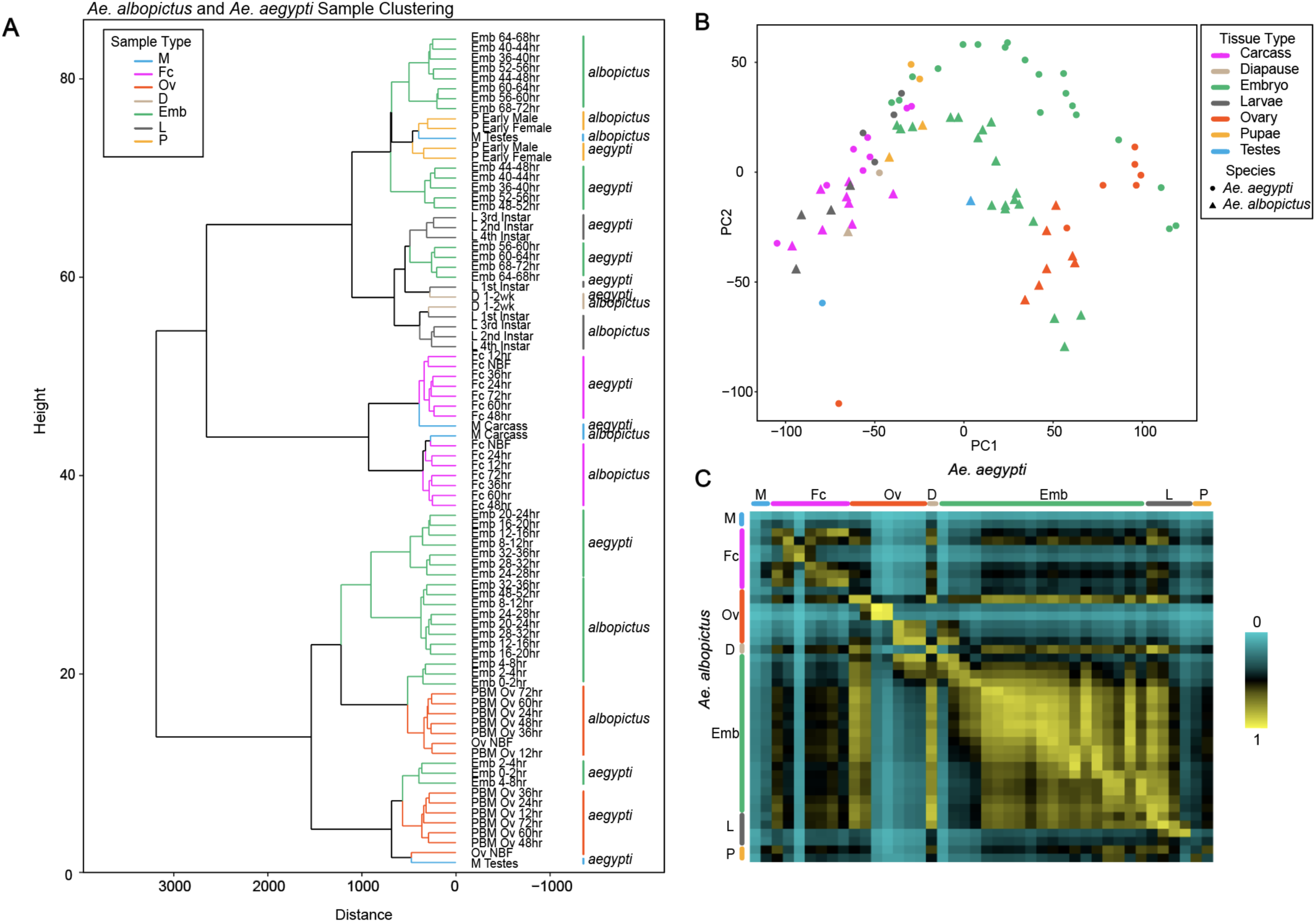
Orthology analysis on *Ae. aegypti* and *Ae. albopictus* samples across corresponding developmental time points. (A) Dendrogram and (B) principal component analysis (PCA) on similar life stage sample types in both species. Both clustering analyses agree with each other indicating the close relationships of similar genes among each developmental time point between species. Interestingly, the *Ae. albopictus* male testes sample clusters distantly from *Ae. aegypti* testes which may indicate a significant difference between the two species. (C) Heat map of calculated Pearson correlations on samples between species. *Ae. albopictus* samples are represented on the vertical axis of the heat map while *Ae. aegypti* samples are represented on the horizontal axis. Life stages are indicated by the similar colored bars for both species.

Ingestion of vertebrate blood is essential for egg maturation by female mosquitoes. In order to see how a blood meal changes gene expression in female ovaries, we performed a differential expression analysis on NBF and PBM ovaries and carcasses (Supplemental Fig. S3; Supplemental Table S9, S10). A distinct pattern emerges when NBF females encounter a blood meal: genes upregulated in NBF samples are downregulated in PBM samples and vice versa (Supplemental Fig. S3). In PBM ovaries samples, ten genes encoding vitelline membrane protein 15a are overexpressed in 12-, 24- and 36hr and correspond to more than 70% of the FPKM expressed in this tissue and time points. In 48hr PBM ovaries, four uncharacterized proteins, two mucins-2, one general odorant-binding protein 45-like, one vitelline membrane protein and one late cornified envelope-like proline-rich protein 1 are overexpressed. The expression of these proteins falls beginning 60hr PBM. In 60- and 72hr PBM ovaries samples only three genes are have FPKM greater than 1,000 and include two uncharacterized loci and a gene that encodes a phosphoglycolate phosphatase 2 (Supplemental Table S9).

We did not observe a significant sex bias between female and male pupae when the early and late pupae stages are compared. However, when we compared the early and late stages of pupae within each sex, we found there was a significant difference between them (Supplemental Fig. S2). Among the top overexpressed genes in female and male late pupae are ejaculatory bulb-specific protein 3, about 10 genes associated with cuticle protein and larval cuticle protein which are important for cuticle formation, carbonic anhydrase 2, zinc metalloproteinase nas-13, and hybrid signal transduction histidine kinase G proteins which are related to protein metabolism. These results indicate that the pupal stage predominantly focused on upregulating genes that will enable adult metamorphosis rather than distinguishing sex.

### Male-specific genes

In performing our global expression analysis, we found that the *Ae. albopictus* male testes sample clustered very distantly when compared to all other time point samples (Fig. 1C, D). To make sure this was not due to an error in sample preparation or single replicate basis, we collected a second male testes replicate and performed a correlation analysis against all samples (Supplemental Table S11). We find that both testes replicates are highly similar (r = 0.903) suggesting that the initial findings are supported. For subsequent analyses, we decided to continue with the first testes replicate. To investigate the distant nature of the male testes sample, we performed a differential expression analysis on male testes and male carcass to find genes that are exclusively expressed in the male. The male germline was compared with female ovaries at NBF and PBM time points. The somatic tissue was also compared to female NBF and PBM carcasses to acquire differentially expressed genes specific to sex. After this initial analysis, we then sorted out the genes which had some expression in other tissue samples. As a result, we found a total of 509 male-specfic genes expressed in the germline and somatic tissues of male mosquitoes (Supplemental Table S12). Of these 509 male-specific genes, 40% of these genes (n = 207) are uncharacterized and function remains to be elucidated. Other genes with relatively high prevalence in the testes include dynein-related genes (6.5%; n = 33), mitochondrial-related genes (3.3%; n = 17), cili- and flagella-associated genes (3%; n = 13), EF-hand related genes (2%; n = 11), myosin-related genes (1%; n = 6), and tubulin-related genes (1.5%; n = 8) (Supplemental Table S12). Within these gene groups, the dynein related genes exhibit variable moderate expression (10 - 111 FPKM), mitochondrial-related genes exhibit relatively high expression (36.6 - 889.9 FPKM), cili- and flagella-associated genes are moderately expressed (26 - 252 FPKM), EF-hand related genes are moderately expressed (50 - 207.6 FPKM), myosin chain-like genes are variably expressed (8 - 158 FPKM), and tubulin genes with relatively high expression (110 - 3074 FPKM) (Supplemental Table S12). Genes with extremely high expression (FPKM > 900) correspond to a few tubulin genes (LOC109407232, LOC109414298, LOC109424041, LOC109426232), several uncharacterized genes, a few cytosol aminopeptidases (LOC109430117, LOC109420901, and LOC109425977), a 36.4kDa proline rich protein (LOC109423840), an arginine kinase (LOC109418450), and a couple keratin-associated proteins (LOC109429278 and LOC109398807) (Supplemental Table S12). It is likely that these genes are crucial for sperm development and management for male reproductive success. Among these exclusive genes we note an interesting observation. Two highly expressed genes, LOC109414298 (Location: NW_017856468.1 340363..342022) and LOC109407232 (Location: NW_017856205.1 (8069187..8070779), found in different locations have 100% amino acid similarity to *Ae. aegypti*’s beta tubulin 2 protein sequence (Smith *et al.* 2007). While the nucleic acid sequence differed between *Ae. aegypti* beta tubulin 2 and LOC109414298 and LOC109407232, their amino acid sequences were highly similar. This suggests that *Ae. albopictus* contains at least two copies of the beta-tubulin 2 gene as paralogs. This finding is supported by a previous finding where twice as many seminal fluid proteins were identified in *Ae. albopictus* compared to *Ae. aegypti* (Degner *et al.* 2019). While we note there is duplication present in our species, we do not yet fully understand how two copies of a gene can affect mosquito biology of *Ae. albopictus*.

### Small RNA pathway protein dynamics

There are three major classes of regulatory small RNAs in animals that include: microRNAs (miRNAs), small interfering RNAs (siRNAs), and Piwi-interacting RNAs (piRNAs). Small RNAs are classified based on their size and interaction with Argonaute proteins. In addition, they are each involved in regulating particular processes in the mosquito. For example, miRNAs are shown to post-transcriptionally regulate transcript levels and translational status mRNA (Lucas *et al.* 2013). In mosquitoes, some miRNAs have been implicated in the regulation and function of blood digestion and ovarian development (Lucas *et al.* 2013). In contrast, the siRNA pathway is responsible for modulating arbovirus replication and can be responsible for transposable element silencing. Finally, the piRNA pathway is suggested to control the remobilization of transposable elements and may take part in antiviral immunity. In *Aedes* mosquitoes, there are 7 PIWI proteins each with a seemingly distinct function (Miesen *et al.* 2015, 2016). To gain a global view of the genes involved in small RNA processing we generated a heatmap to visualize their expression across development (Supplemental Fig. S4; Supplemental Table S13). Increased expression of two piRNA genes, Mael-B (AALF019672) and Gstf (arx) (AALF023639), are apparent in NBF and PBM ovaries (Supplemental Fig. S4). Interestingly, Twin (AALF018294), a CCR4 deadenylase, is highly expressed when the embryos undergo diapause. In *Drosophila*, this gene is shown to promote the decay of specific mRNAs in the early embryo (Rouget *et al.* 2010). In *Ae. albopictus*, it may serve as a mechanism to ensure diapause is transcriptionally arrested until there is an environmental signal that induces diapause termination. In contrast to all the stages, the male and female NBF and PBM germline highly expresses PIWI genes 2 and 3. While there are several functions for the use of piRNAs in the insect germline, it is likely this observation suggests that the PIWI pathway is involved in silencing retrotransposons (Kalmykova *et al.* 2005; Wang and Elgin 2011). PIWI3 was found to be downregulated in male testes, later PBM ovaries, and in the early embryo (0-1hr and 0-4hr). While there is still not much known about each particular PIWI protein, a study found that PIWI3 may be associated with viral dissemination in mosquitoes (Wang *et al.* 2018).

### Comparison between *Ae. albopictus* and *Ae. aegypti* development transcriptomes

#### Transcriptome overview

We next sought to determine the similarities and differences between the developmental transcriptomes of *Ae. aegypti* and *Ae. albopictus* mosquitoes. To establish orthologues, we performed BLAST searches on the proteomes of both species and identified best reciprocal hits ensuring one-to-one relationship between genes in the two species resulting in 10,696 orthologous pairs representing 54.51% and 27.63% of genes encoded by *Ae. aegypti* and *Ae. albopictus* genomes, respectively (Supplemental Table S14 for best reciprocal hits). Using previously published *Ae. aegypti* developmental transcriptome data (Akbari *et al.* 2013a), we conducted a set of analyses comparing gene expression levels between the two species’ developmental stages with the aim to gain insight into possible differences in their biology (Supplemental Table S1). On average, 57.6 % (range: 23.3% to 72.9%) and 34.6% (range: 29.1% to 39.4%) of RNA-seq reads from *Ae. aegypti* and *Ae. albopictus* datasets were mapped to orthologs, respectively (Supplemental Table S15). Sample clustering and PCA analysis revealed that the majority of the corresponding sample timepoints and tissues between both species display similar overall expression patterns and are found adjacent to each other (Fig. 3 A, B). A notable exception is the testes sample pair, which cluster far apart presumably reflecting significantly different gene expression programs. Finally, we calculated Pearson correlations between FPKM values of the corresponding samples and then plotted them on a heatmap to confirm similarities between embryonic samples (Fig. 3C, Supplemental Table S16). Embryonic samples between species have higher correlations indicating that the genes involved in this developmental stage are very similar or shared. This is also true for all other tissues/time points, however, with the exception of the male testes, consistent with the results of clustering analysis (Fig. 3C; Fig. S5; Supplemental Table S16).

We next performed a differential expression analysis between *Ae. albopictus* and *Ae. aegypti* samples at each developmental time point to gain some insight into the differences of orthologous genes expressed (Supplemental Table S17). Supplemental Figure S6 depicts the number of orthologous genes differentially expressed (with a *padj* < 0.1) between both species at each developmental stage. Across the whole cycle of development, the male testes, NBF female ovaries, 36hr PBM ovaries, 48-52hr and 64-68hr embryo, 4^th^ instar larvae, and pupae stages have distinct patterns of expression between *Ae. albopictus* and *Ae. aegypti*. In the sections below, we delve further into the identity and function of these genes.

#### Male Germline

Based on our initial analyses of the *Ae. albopictus* samples, the male testes sample depicted major differences when compared to all stages (Fig. 1). It was highly uncorrelated to other stages within this species and displayed a uniqueness that needed to be further investigated. When the testes sample was compared to *Ae. aegypti*, we found that more genes were upregulated in *Ae. albopictus* testes as compared to *Ae. aegypti* testes (172 versus 97 genes, respectively) (Supplemental Fig. S6, S7; Supplemental Table S17). The majority of these orthologous genes in *Ae. albopictus* (n = 69) are uncharacterized and their function remains unknown (Supplemental Table S17). It would be interesting to determine the function and significance these genes contribute to the male germline development and perhaps fertility. Several annotated genes correspond to flagellar structure including microtubules, dynein, ciliar components, and proteins associated with mitochondrial derivatives (Supplemental Table S17). Other genes include testis-specific protein kinases, histone genes, and several genes involved in DNA-binding transcription factor activity, presumably to regulate transcription in spermatogenesis. Several serine/threonine-protein kinases are expressed and are involved in the control of many physiological processes, like flagellar motility and muscle contraction (Cohen 1997). Cytosol aminopeptidase was also upregulated in *Ae. albopictus*. This gene is among one of the top highly expressed male-specifc genes in our *Ae. albopictus* testes analyses, with multiple copies present, and is known to be one of the most abundant sperm proteins in *Ae. aegypti* seminal fluid (Degner *et al.* 2019). In *Ae. aegypti*, relatively fewer genes are upregulated and have about 25 uncharacterized genes (Supplemental Table S17). In *Ae. aegypti* testes, odorant binding proteins, carboxypeptidases, trypsins, and hormone-related genes are highly expressed compared to *Ae. albopictus*. These up-regulated hormone-related genes (juvenile hormone esterase-like, venom carboxylesterase-6-like, and juvenile hormone esterase-like) are shown to be down-regulated in 24 hr post mated *Ae. aegypti* females and play a major role during female reproduction (Alfonso-Parra *et al.* 2016). In *Anopheles gambiae* males, steroid hormones are generated and stored in the male accessory glands and are transferred during mating (Pondeville *et al.* 2008). It is likely that this physiological process is implicated in the reproductive success of *An. gambiae* mosquitoes and may apply to *Ae. aegypti* but not in *Ae. albopictus*. However, male *Ae. albopictus* may have evolved different sexual strategies to ensure reproductive success (Oliva *et al.* 2013; Bargielowski and Lounibos 2014).

#### Female Germline (NBF and PBM 36hr ovaries)

Several studies have been performed on the blood feeding behavior of *Ae. albopictus* and *Ae. aegypti* female mosquitoes. While helpful in understanding the female behavioral differences between the two species, it is important to understand the underlying transcriptional basis that contribute to these differences. In NBF ovaries, *Ae. aegypti* has 205 significantly upregulated genes compared to *Ae. albopictus*, who has 9 (Supplemental Supplemental Figs. S5, S6; Supplemental Table S17). The majority of the upregulated genes within *Ae. aegypti* NBF ovaries are uncharacterized and do not associate with any GO enrichment term, however, those that do deal mostly with DNA binding, zinc ion binding, protein binding, translation, cell-redox homeostasis, and regulation of transcription (Supplemental Table S17). In *Ae. albopictus*, GO enrichment terms associated with ATP binding, protein binding, translation, methylation, and metabolic processes are prevalent in this sample (Supplemental Fig. S17). Genes overrepresented are related to the mitochondria including NADH dehydrogenase, cytochrome c, and ATP synthase; and ribosomal genes encoding 60S and 40S ribosomal proteins important for translation (Supplemental Table S17). The observation that *Ae. aegypti* NBF ovaries have more upregulated genes than *Ae. albopictus* supports a previous finding where *Ae. aegypti* exhibits fast evolution of ovary-biased genes compared to *Drosophila melanogaster* and *Ae. albopictus* (Whittle and Extavour 2017).

When female mosquitoes are blood fed, a wave of gene expression is initiated to metabolize proteins and nutrients from the blood meal to provide nutrients for her eggs (Zhou *et al.* 2007). We wondered how different gene expression programs are between *Ae. albopictus* and *Ae. aegypti* given that these two species have somewhat different feeding behaviors. According to our DESeq analysis, 12hr PBM ovaries, 24hr PBM carcass, and 72hr PBM carcass samples had no significant differentially expressed genes indicating similar gene expression programs in the two species (Supplemental Fig. S7). The sample that had the most differentially expressed genes in *Ae. albopictus* was the 36hr PBM ovaries, where 48 orthologous genes highly expressed compared to 9 in *Ae. aegypti* (Supplemental Fig. S7; Supplemental Table S17). In *Ae. aegypti*, ecdysone-induced protein, histone H2A-like, heat shock 70 protein, chloride channel proteins and others are upregulated (Supplemental Fig. S7; Supplemental Table S17). Only eight genes in *Ae. albopictus* were found to be uncharacterized while genes coding for major royal jelly proteins, odorant-binding proteins, mucin, and vitellogenin are prevalent in this species. The vitellogenin and odorant binding protein are consistent with their involvement in the maturation of vitellogenic oocytes and eggshell formation (Costa-da-Silva *et al.* 2013). It is interesting to note that royal jelly proteins are highly expressed in the developing zygote in female *Ae. aegypti* mosquito ovaries.

#### Late embryonic stages (48-52hr and 64-68hr)

Only two timepoints during embryogenesis significantly differ between both species. In both stages, 48-52-hr and 64-68hr, *Ae. aegypti* exhibits a very high number of upregulated genes (247 genes and 350 genes, respectively) (Supplemental Fig. S8; Supplemental Table S17). About 62 uncharacterized genes are significantly differentially expressed in *Ae. aegypti* 48-52hr embryos. Several genes coding for cuticle proteins are enriched in this late embryonic stage and suggest the importance of embryonic desiccation resistance. Mucin-like protein coding genes were also enriched and may reflect the protective function on developing organ epithelial surfaces of the embryo (Syed *et al.* 2008). In *Ae. albopictus* several genes related to mitochondrial function are upregulated compared to *Ae. aegypti*. Because mitochondria are key organelles in obtaining energy from nutrients by means of oxidative phosphorylation, it is possible that late embryogenesis in *Ae. albopictus* requires high energy input for cellular processes than *in Ae. aegypti*. During the 64-68hr embryonic stage, *Ae. albopictus* only contains 39 differentially expressed genes compared to *Ae. aegypti* (Supplemental Table S17). About 13 genes are uncharacterized and genes such as odorant binding protein 56d, cuticle protein, mucin, and mitochondrial translocase are present at this stage. In *Ae. aegypti*, several genes related to cuticle (n = 66) and trypsins (n = 25) are enriched.

#### Larval stages (4^th^ instar) and pupal stages (male and female)

The larval and pupal stages represent important aquatic stages of the mosquito life cycle. In aquatic habitats, mosquitoes must develop and survive to reach the adult stage, the stage that will ultimately have epidemiological consequences. Both *Ae. albopictus* and *Ae. aegypti* are found to survive better in urbanized areas (Li *et al.* 2014) and can co-occur in the same breeding sites. In areas where both species are present, it has been observed that *Ae. albopictus* can successfully displace *Ae. aegypti* populations. One hypothesis suggests that *Ae. albopictus* is a superior larval competitor and can withstand larvae starvation longer (Juliano 2009). To get an overview of the molecular differences between these two species, we performed a DESeq2 analysis on larvae samples. In our analyses we find differentially expressed genes in all instar stages except for the second (Supplemental Fig. S9; Supplemental Table S17). Only 18 significantly upregulated orthologs were present in *Ae. albopictu*s and include 5 uncharacterized loci, a couple of mitochondrial-related genes, and a pupal cuticle protein. In contrast, *Ae. aegypti* has 48 upregulated orthologs and include genes enriched for uncharacterized loci and cuticle proteins (Supplemental Table S17).

In *Ae. albopictus* male pupae, only 7 genes were found to be significantly upregulated as compared to *Ae. aegypti* who has 65 orthologous genes upregulated (Supplemental Fig. S6; Supplemental Table S17). Among the seven, a protease known as neprilysin was found to be highly expressed in *Ae. albopictus* male pupae. Neprilysins play an important role in turning off peptide signaling events at the surface of the cell (Turner *et al.* 2001). In *D. melanogaster*, neprilysins were reported to be expressed in the malpighian tubules and testes suggesting roles for peptidase activity in the excretory systems and in spermatogenesis (Thomas *et al.* 2005). In addition, a mitochondrial dimethyladenosine transferase is expressed and is essential for the maturation and maintenance of the mitochondrial ribosome (Raimundo 2014). The presence of lysosomal alpha-mannosidase gene suggests male pupae place importance of metabolizing mannose-containing oligosaccharides (Paciotti *et al.* 2017). The lysosomal catabolism of glycoproteins is crucial for maintaining cellular homeostasis, however, its roles for male pupae development remain elusive. Of the 60 significantly differentially expressed orthologs in *Ae. aegypti*, 30 genes correspond to cuticle proteins, 3 correspond to mitochondria-related genes, and 2 correspond to odorant-binding proteins (Supplemental Table S17). In female pupae samples, 20 and 91 orthologous genes were significantly upregulated in *Ae. albopictus* and *Ae. aegypti*, respectively. Only 6 cuticle-related genes, a neprilysin, and a couple of mitochondrial genes were up-regulated in *Ae. albopictus*. Out of 91 significantly differentially expressed genes in *Ae. aegypti* female pupae, 26 of them correspond to cuticle genes, 4 are mitochondrial-related, 19 are uncharacterized, and two chorion-related genes (Supplemental Fig. S6; Supplemental Table S17). Chorion protein and chorion peroxidases are usually highly expressed in early embryonic stages and in adult female oocytes, however it is likely that their expression in the pupae reflect developing female genitalia this early in the life cycle (Chambers 2005).

## Discussion

*Ae. aegypti* and *Ae. albopictus* are medically important mosquitoes with the capacity to transmit a variety of human pathogens. Taking into account that these mosquitoes are vectors of dengue, Zika and chikungunya viruses, there is a potential risk of increasing the incidence of these diseases. Although *Ae. aegypti* is considered a main vector for these viruses, *Ae. albopictus* is emerging as another important vector. In order to contribute to the knowledge of the biological development of *Ae. albopictus*, we analyzed the whole transcriptome at different stages of the life cycle. The observations presented here should reflect a comprehensive snapshot of the *Ae. albopictus* developmental transcriptome. Our results provide confirmation for up to 95% (36,347/38,261) of previously annotated AALF genes.

The mosquito’s life cycle can be divided into 4 major phases: the maternal to zygotic transition (ovary to embryos 0-8hr); embryogenesis (from 8- to 72hr), diapause (0-1wk and 2-3wks); larvae (1-4 instars) to pupae (early and late); and adults (male and female). These crucial transitions of the mosquito life cycle share many genes whose expression shows little difference between each time point. Our cluster analysis and characterization of developmental and sex-specific genes identified a number of patterns of co-regulated gene expression. We see significant differential expression in male and female germline that gives us insights into the reproductive biology of *Ae. albopictus*. Identification of loci involved in the blood meal program of the ovaries will be of interest to understand the regulation of ovarian development. In this study, we were able to depict the dynamics of genes involved in small RNA production (siRNAs, miRNAs, and piRNAs) across all development. Small RNAs in mosquitoes are known to partake in many important roles in cell development, response to stress, infection, and the silencing of transposable elements (Lucas *et al.* 2013). While we did not characterize small RNAs, our analysis on genes involved in small RNA production gives us insights into the roles of the small RNA pathways in *Ae. albopictus*. It would be interesting to investigate the small RNA profiles as it pertains to our results.

In our transcriptomic analysis of mosquito tissues, we discovered that the male testes showed a distinct gene expression profile that differentiated it from other tissues. Upon closer inspection, we identified 509 male-biased genes that were expressed in male testes, carcass, and male pupae. The highest expressing genes correspond to several uncharacterized loci, cytosol aminopeptidases, tubulins, and a 36.4kDa proline rich protein. It would be interesting to see what functions and processes the uncharacterized genes are involved in. It is likely they may be involved in spermatogenesis, seminal fluid production, or a mating induced response. Perhaps these highly expressed genes have important roles in spermatogenesis and in the management and production of seminal fluid proteins to enable the reproductive success of *Ae. albopictus* males. The highly divergent gene expression in the testes may suggest male-specific genes in this species are evolving rapidly (Swanson and Vacquier 2002). *Ae. albopictus* males exhibit an interesting mating strategy that involves male mosquitoes mating with multiple females in succession despite the availability of sperm in their testes (Oliva *et al.* 2013). Another strategy involves transferring male accessory gland secretions with sperm into the female to prevent further insemination. This is very similar to the mating plug seen in *Anopheline* species (Giglioli 1964). In the wild, *Ae. albopictus* mosquitoes are shown to displace *Ae. aegypti* populations in areas where they co-occur (Muzari *et al.* 2019). Currently, it is hypothesized that *Ae. albopictus* is able to do this because it is a superior larval competitor, however, a recent study suggested that competitive displacement is due to *Ae. albopictus* males mating with *Ae. aegypti* females results in female mating refractoriness (Tripet *et al.* 2011). This mating interference is known as ‘satyrization’ and has been suggested to be a form of adaptation favoring the invasive success of *Ae. albopictus* (Lounibos and Kramer 2016). In our search to understand what are the highly expressed genes in male testes, we found at least one beta-tubulin-2 gene that was duplicated. This is not surprising considering the *Ae. albopictus* genome is highly duplicated, however, it is unclear to what extent a duplication of male-biased genes can contribute to the male reproductive success of these mosquitoes. It is possible that in this species, the testes-biased genes can exhibit rapid evolution contrary to *Ae. aegypti* which experiences decelerated rates of evolution in the testes (Whittle and Extavour 2017).

The comparative *Aedes* developmental transcriptomics approach enabled us to observe differences in developmental life stages. Specifically, we find that there are seven sample/tissue time points including male testes, NBF ovaries, 36hr PBM ovaries, two late embryonic stages (48-62hr and 64-68hr), 4^th^ instar larvae, and pupal stage that have considerably different gene expression dynamics (Supplemental Fig. S6; Supplemental Table S17). Generally, *Ae. aegypti* displays a higher number of significantly upregulated genes in all these stages with the exception of the male testes. The number of upregulated genes in *Ae. aegypti* NBF ovaries are significantly higher than *Ae. albopictus*. Genes overrepresented the NBF ovaries include those with mitochondrial ATP synthesis function and translation. Because mitochondrial genomes (mtDNA) are strictly maternally transmitted they can only make a direct evolutionary response through females (Innocenti *et al.* 2011). Perhaps selection in mtDNA that contributes to sex-specific traits (sexually dimorphic or sexually antagonistic) may be a mechanism for the *Ae. aegypti* ovaries to evolve rapidly (Innocenti *et al.* 2011; Whittle and Extavour 2017). Genes in the reproductive tissues contributing to the sexual and reproductive functions are often rapidly evolving (Jagadeeshan and Singh 2005). The observation that *Ae. albopictus* has a higher number of upregulated genes in the testes compared to *Ae. aegypti* suggests that the male germline may experience rapid evolution of male-bias genes in this species. In 36hr PBM ovaries, we find two genes related to major royal jelly protein family significantly differentially expressed in *Ae. aegypti* samples. Royal jelly has been implicated in having a central role in honeybee (*Apis mellifera*) development and genes encoding major royal jelly proteins (5 in total) share a common evolutionary origin with the yellow pigmentation gene of *Drosophila melanogaster* (Albert *et al.* 1999). While it is not known for mosquitoes to utilize royal jelly, it has been shown in *Drosophila melanogaster* that a diet of major royal jelly proteins have resulted in increases in lifespan, feeding, and fecundity (Xin *et al.* 2016). It remains to be determined if the two major royal jelly proteins (1 and 5) present in *Ae. albopictus* 36hr PBM ovaries are actually functional. If this were the case, it may explain an important difference between the two species.

In addition to providing a tool for basic molecular research on *Ae. albopictus*, the developmental transcriptome will facilitate the development of transgenesis-based control of vector populations. Regulatory elements that direct expression of transgenes in germline specific tissues will be useful for the development of gene drive mechanisms for spreading a desired trait in a mosquito population (Sieglaff *et al.* 2009; Akbari *et al.* 2014b). In *Aedes aegypti*, several regulatory elements able to drive gene expression in a tissue- and temporal-specific manner have been identified through extensive study (Akbari *et al.* 2013a) and transgenesis (Coates *et al.* 1999; Kokoza *et al.* 2000; Moreira *et al.* 2000; Smith *et al.* 2007). Future functional characterization of uncharacterized genes and regulatory elements may lead to the development of innovative genetic population control technologies such as precision guided sterile males (Kandul *et al.* 2019b), and gene drive systems (Akbari *et al.* 2013b, 2014a; Champer *et al.* 2016; Buchman *et al.* 2018b, 2018a; Kandul *et al.* 2019a, 2019b; Li *et al.* 2019) which can be linked to anti-pathogen effectors (Buchman *et al.* 2019a, 2019b) potentially providing paradigm-shifting technologies to control this worldwide human disease vector. Overall, our results provide a comprehensive snapshot of gene expression dynamics in the development of *Ae. albopictus* mosquitoes. The comparative analysis performed between *Ae. albopictus* and *Ae. aegypti* will be helpful in facilitating future comparative biological studies to understand the molecular basis of their differences.

## Methods

### Mosquito strain

Mosquitoes used for RNA extraction were from wildtype *Ae. albopictus* which originated from San Gabriel Valley, located in the Los Angeles County, CA. Mosquito eggs were collected from oviposition traps set by the San Gabriel Valley Mosquito Control district. Collected eggs were from sites were *Ae. albopictus* was known to circulate. These eggs were then hatched, checked for the characteristic stripe of *Ae. albopictus,* and reared for 10 generations before performing collection experiments. Mosquitoes were maintained in insectary facility with a relative humidity of 70-80%, maintained at 28C, and with a 12-hr/12hr light/dark cycle. Larvae were fed with ground fish food (TetraMin Tropical Flakes, Tetra Werke, Melle, Germany) and sex separated as pupae. Adults were maintained and fed with an aqueous solution of 10% sucrose. Females were blood-fed 3-5 days after eclosion on anesthetized mice. All animals were treated according to the Guide for the Care and Use of Laboratory Animals as recommended by the National Institutes of Health.

### Total RNA isolation and RNA sequencing

Samples were flash-frozen at specific time points, and total RNA was extracted using the Ambion mirVana mRNA isolation kit (Ambion/Applied Biosystems, Austin, TX). The total RNA for the second testes replicate was extracted using the Qiagen RNeasy Mini kit (Qiagen, Germantown, MD). All sample collections were staged in the incubator at a relative humidity of 70–80%, 28° with a 12-hr/12-hr light cycle until the desired time point was reached. Samples were then immediately flash frozen. The adult NBF carcass was processed at 3 d after eclosion, and the adult male carcass and testes were processed at 4 d after eclosion. After extraction, RNA was treated with Ambion Turbo DNase (Ambion/Applied Biosystems, Austin, TX). RNA integrity was assessed using RNA 6000 Pico Kit for Bioanalyzer (Agilent Technologies #5067-1513). RNA-seq libraries were constructed using NEBNext Ultra II RNA Library Prep Kit for Illumina (NEB #E7770) following manufacturer’s instructions. Briefly, mRNA isolated from ∼1 μg of total RNA was fragmented to the average size of 200 nt by incubating at 94 °C for 15 min in first strand buffer, cDNA was synthesized using random primers and ProtoScript II Reverse Transcriptase followed by second strand synthesis using NEB Second Strand Synthesis Enzyme Mix. Resulting DNA fragments were end-repaired, dA tailed and ligated to NEBNext hairpin adaptors (NEB #E7335). After ligation, adaptors were converted to the ‘Y’ shape by treating with USER enzyme and DNA fragments were size selected using Agencourt AMPure XP beads (Beckman Coulter #A63880) to generate fragment sizes between 250 and 350 bp. Adaptor-ligated DNA was PCR amplified followed by AMPure XP bead clean up. Libraries were quantified with Qubit dsDNA HS Kit (ThermoFisher Scientific #Q32854) and the size distribution was confirmed with High Sensitivity DNA Kit for Bioanalyzer (Agilent Technologies #5067-4626). Libraries were sequenced on Illumina HiSeq2500 in single read mode with the read length of 50 nt following manufacturer’s instructions. Base calls were performed with RTA 1.18.64 followed by conversion to FASTQ with bcl2fastq 1.8.4.

### Poly(A+) read alignment and quantification

Reads from RNA-seq libraries were aligned to the *Ae. albopictus* genome (canu_80X_arrow2.2, GCA_001876365.2) using STAR aligner (Dobin *et al.* 2013). Gene models were downloaded from NCBI (ftp://ftp.ncbi.nlm.nih.gov/genomes/all/GCF/001/876/365/GCF_001876365.2_canu_80X_arrow2.2/GCF_00187636.5.2_canu_80X_arrow2.2_genomic.gff.gz) and quantified with featureCounts (Liao *et al.* 2014). TPM (Transcripts Per Million) and FPKM (Fragments Per Kilobase Million) values were calculated from count data using Perl scripts. All Sequencing data has been made publically available at NCBI Sequence Read Archive (SRA) under Accession number (will provide upon publication).

### Clustering and Gene Ontology (GO) analysis

TPM values produced by featureCounts for 47 RNA-seq libraries were clustered using Mfuzz R software package (Kumar and E Futschik 2007). Mfuzz uses fuzzy c-means algorithm to perform soft clustering, which allows cluster overlap and has been demonstrated to perform favorably on gene expression data. The resulting clusters were analyzed for overrepresentation of GO terms using a hypergeometric test implemented using the GOstats R software package (Falcon and Gentleman 2007). Pfam domains for the *Ae. Albopictus* gene set were identified by running hmmscan (Finn *et al.* 2011) and associated GO terms were added using pfam2go mapping downloaded from the Gene Ontology Consortium (The Gene Ontology Consortium and The Gene Ontology Consortium 2019). Hypergeometric tests were performed separately for biological process, molecular function, and cellular component ontologies. Sample dendrograms and PCA plots were generated in R and plotted with ggdendro and ggplot2 packages. Up and down regulated genes between developmental stages were identified with DESeq2 R package (Love *et al.* 2014).

### Comparative analysis between *Ae. albopictus* and *Ae. aegypti* transcriptomes

Ortholog pairs between *Ae. albopictus* and *Ae. aegypti* were assembled by running blastp on protein sets (GCF_001876365.2_canu_80X_arrow2.2_protein.faa and GCF_002204515.2_AaegL5.0_protein.faa) for the two species against each other and identifying best reciprocal hits resulting in 10,696 orthologous gene pairs. That represents 54.51% and 27.63% of genes encoded by *Ae. aegypti* and *Ae. albopictus* genomes, respectively. The lower percent of the *Ae. albopictus* gene set representation is likely the result of extensive duplication known to exist in the current version of *Ae. albopictus* genome. Ortholog quantification was performed with featureCounts and genes differentially expressed between the two species for each developmental stage were identified using DESeq2.

## Acknowledgements

This work was supported by funding from generous UCSD lab startup funds awarded to O.S.A. We thank Vijaya Kumar for help with library preparations. Transcriptome sequencing was performed at the Millard and Muriel Jacobs Genetics and Genomics Laboratory at the California Institute of Technology. We also thank Dr. J. Wakoli Wekesa and the San Gabriel Valley Mosquito and Vector Control District for providing us with the *Ae. albopictus* mosquito eggs used in this study.

## Author Contributions

O.S.A., and I.A. conceptualized the study. S.G. performed all sample preparations. I.A, S.G., and S.C.M performed the various bioinformatic analysis in the study. All authors contributed to the writing, analyzed the data, and approved the final manuscript.

## Supplementary Appendix Contents

❖ SI Appendix Figures
  ➢ **SI Appendix Fig. S1:** Read mapping statistics and read count data across all developmental stages
  ➢ **SI Appendix Fig. S2:** Volcano plots visualize the fold-difference in individual gene expression between sex-specific tissue samples of *Ae. albopictus*
  ➢ **SI Appendix Fig. S3:** Hierarchical heat map profile of the top 500 differentially expressed genes in the germline and somatic female tissues compromising of ovary or carcass tissues from sugar fed (NBF) and 12hr, 24hr, 36hr, 48hr, 60hr, and 72hr post blood fed female mosquitoes.
  ➢ **SI Appendix Fig. S4:** Expression dynamics of protein genes involved in miRNA, siRNA and piRNA pathways.
  ➢ **SI Appendix Fig. S5:** Sample correlations of each developmental time point between *Ae. aegypti* and *Ae. albopictus*
  ➢ **SI Appendix Fig. S6:** Number of upregulated genes in *Ae. aegypti* and *Ae. albopictus* samples across corresponding developmental time points
  ➢ **SI Appendix Fig. S7:** MA plots of upregulated genes in sex-specific tissues
  ➢ **SI Appendix Fig. S8:** MA plots of upregulated genes in the PBM ovaries and carcass samples of *Ae. albopictus* compared to *Ae. aegypti*
  ➢ **SI Appendix Fig. S9:** MA plots of upregulated genes in embryogenesis of *Ae. albopictus* compared to *Ae. aegypti*
  ➢ **SI Appendix Fig. S10:** MA plots of upregulated genes in the larval instars and diapause samples of *Ae. albopictus* compared to *Ae. aegypti*
❖ SI Appendix Tables
  ➢ **SI Appendix Table S1:** Summary of sequenced datasets
  ➢ **SI Appendix Table S2:** Read mapping analysis of *Ae. albopictus* transcripts
  ➢ **SI Appendix Table S3:** Gene descriptions of *Ae. albopictus* loci
  ➢ **SI Appendix Table S4:** Complete developmental transcriptome transcripts of *Ae. albopictus*.
  ➢ **SI Appendix Table S5:** AALF genes mFuzz cluster membership
  ➢ **SI Appendix Table S6:** Gene ontology enrichment analysis of *Ae. Albopictus* clusters 1 through 20
  ➢ **SI Appendix Table S7:** Early zygotic genes
  ➢ **SI Appendix Table S8:** Strictly maternal genes
  ➢ **SI Appendix Table S9:** Female-specific genes
  ➢ **SI Appendix Table S10:** Ovary-specific genes
  ➢ **SI Appendix Table S11:** Pearson correlation matrix with second male testes replicate
  ➢ **SI Appendix Table S12:** Male-specific genes
  ➢ **SI Appendix Table S13:** Genes involved in small RNA production
  ➢ Read mapping analysis of *Ae. albopictus* transcripts to *Ae. aegypti* genome assembly
  ➢ **SI Appendix Table S14:** Best reciprocal hits orthology
  ➢ **SI Appendix Table S15:** Count summary of albopictus reads mapping to orthologs
  ➢ **SI Appendix Table S16:** Pearson correlation values of *Ae. aegypti* and *Ae. albopictus* at each developmental stage
  ➢ **SI Appendix Table S17:** Upregulated genes in *Ae. aegypti* and *Ae. Albopictus* expressed orthologs

---

**Supplemental Figure 1.**
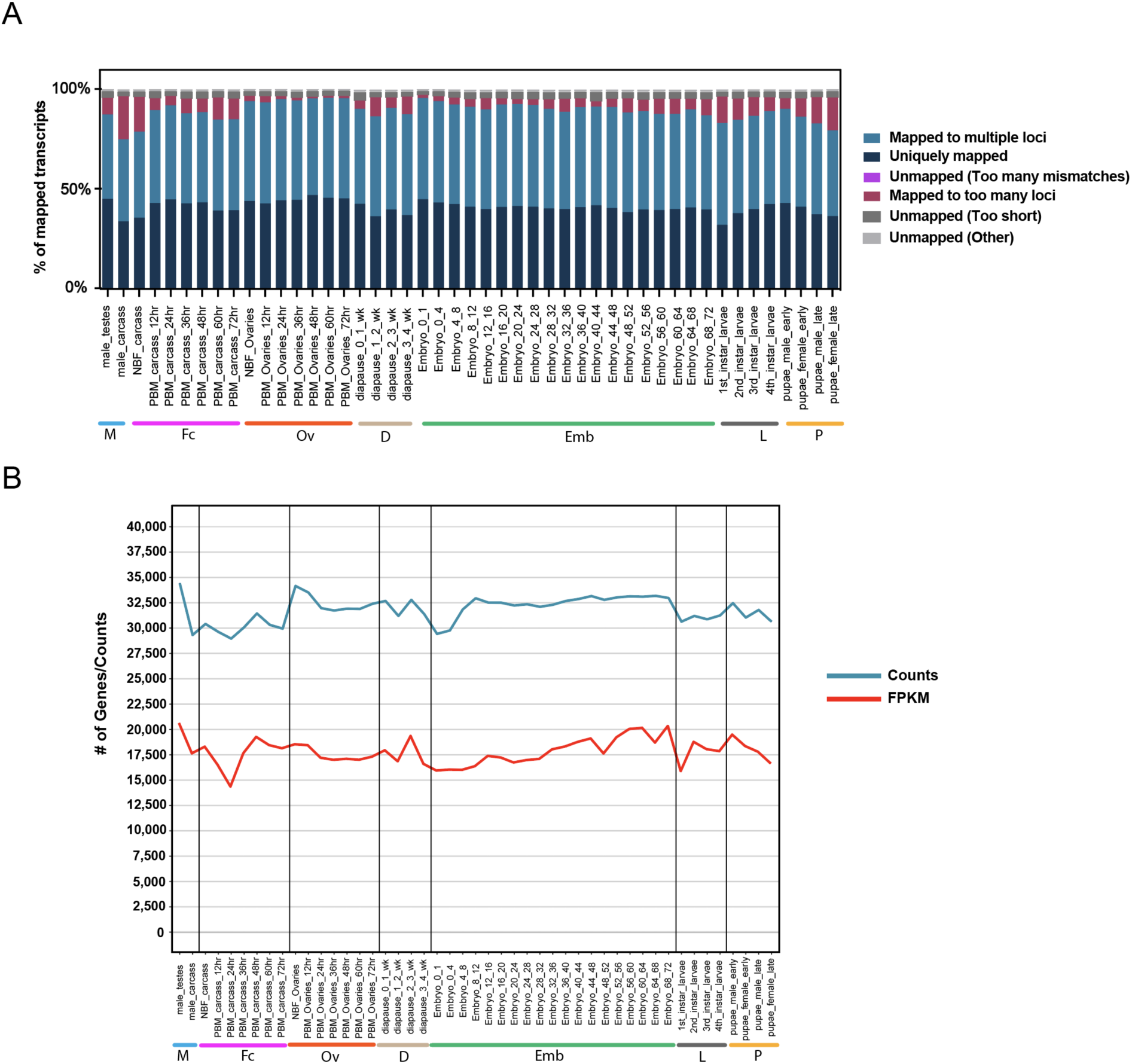
R**e**ad **mapping statistics and read count data across all developmental stages.** (A) Read mapping analysis of *Ae. albopictus* developmental samples. The distribution reflects percentage of fragments mapped to multiple loci (blue), uniquely mapped fragments (dark blue), unmapped fragments with too many mismatches (purple-pink), fragments mapped to too many loci (maroon), unmapped fragments that are too short (dark gray) or other (gray). (B) The number of albopictus genes expressed at FPKM > 1 (red) and with non-zero count values (blue) were plotted across all developmental timepoints. Mapping statistics can be found in Supplemental Table S2.

**Supplemental Figure 2.**
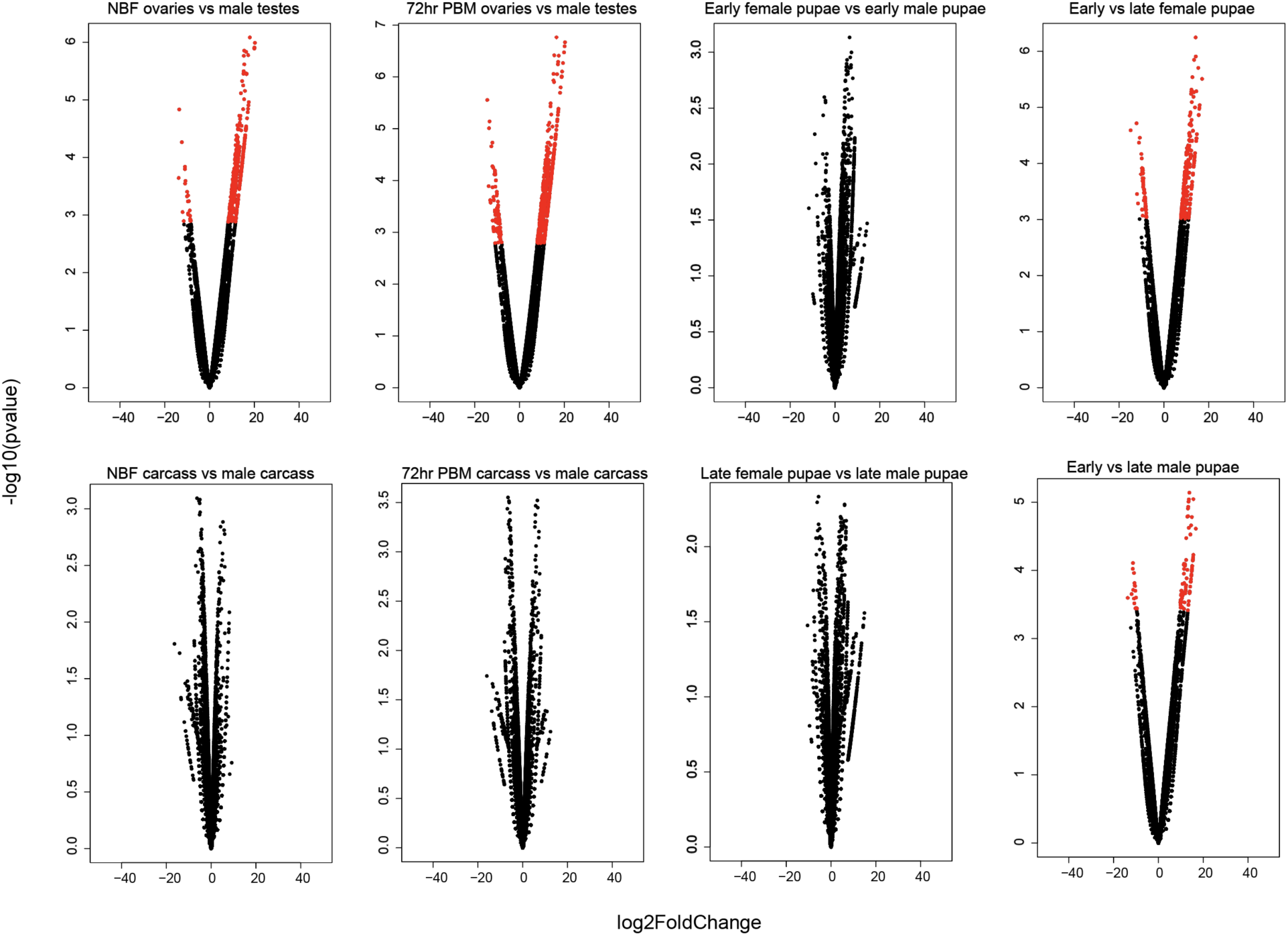
Volcano plots visualize the fold-difference in individual gene expression between sex-specific tissue samples of *Ae. albopictus*. Each gene is represented by a single point. The x-axis displays the log2-transformed signal intensity differences between tissues; positive values represent overexpression in the male tissues with the exception of the right-most top plot. Negative values represent overexpression in female tissues with the exception of the right-most bottom plot. The y-axis displays log10-transformed *P*-values of differential gene expression. Genes that are colored red if they are significantly differentially expressed between the two tissues.

**Supplemental figure S3.**
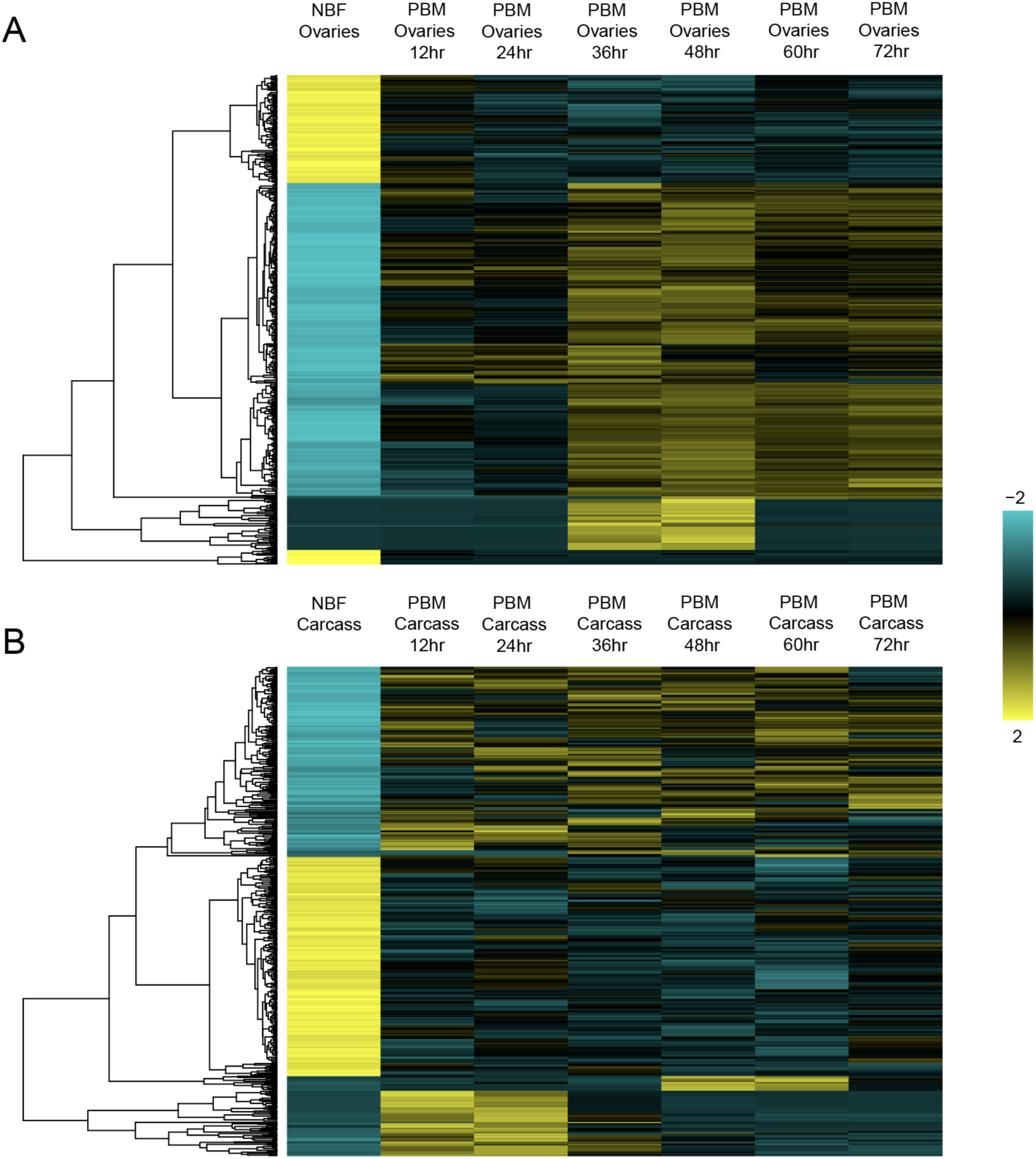
Hierarchical heat map profile of the top 500 differentially expressed genes in the germline and somatic female tissues compromising of ovary or carcass tissues from sugar fed (NBF) and 12-, 24-, 36-, 48-, 60-, and 72hr post blood fed female mosquitoes. There is clear transitioning between samples and different levels of expression of genes within each sample/tissue. (A) In the NBF ovary sample, many genes that are found to be differentially expressed in the other stages are in the lower spectrum, indicating they have either low to no expression. Once the female feeds, there is a clear increase in gene expression. (B) In somatic cells, there is a subset of genes in the NBF female that are highly expressed but are turned down after a blood meal. Each gene is represented by a single row of colored boxes; each time point is represented as a single column of colored boxes. The scale bar indicates the count z-scores.

**Supplementary Figure 4.**
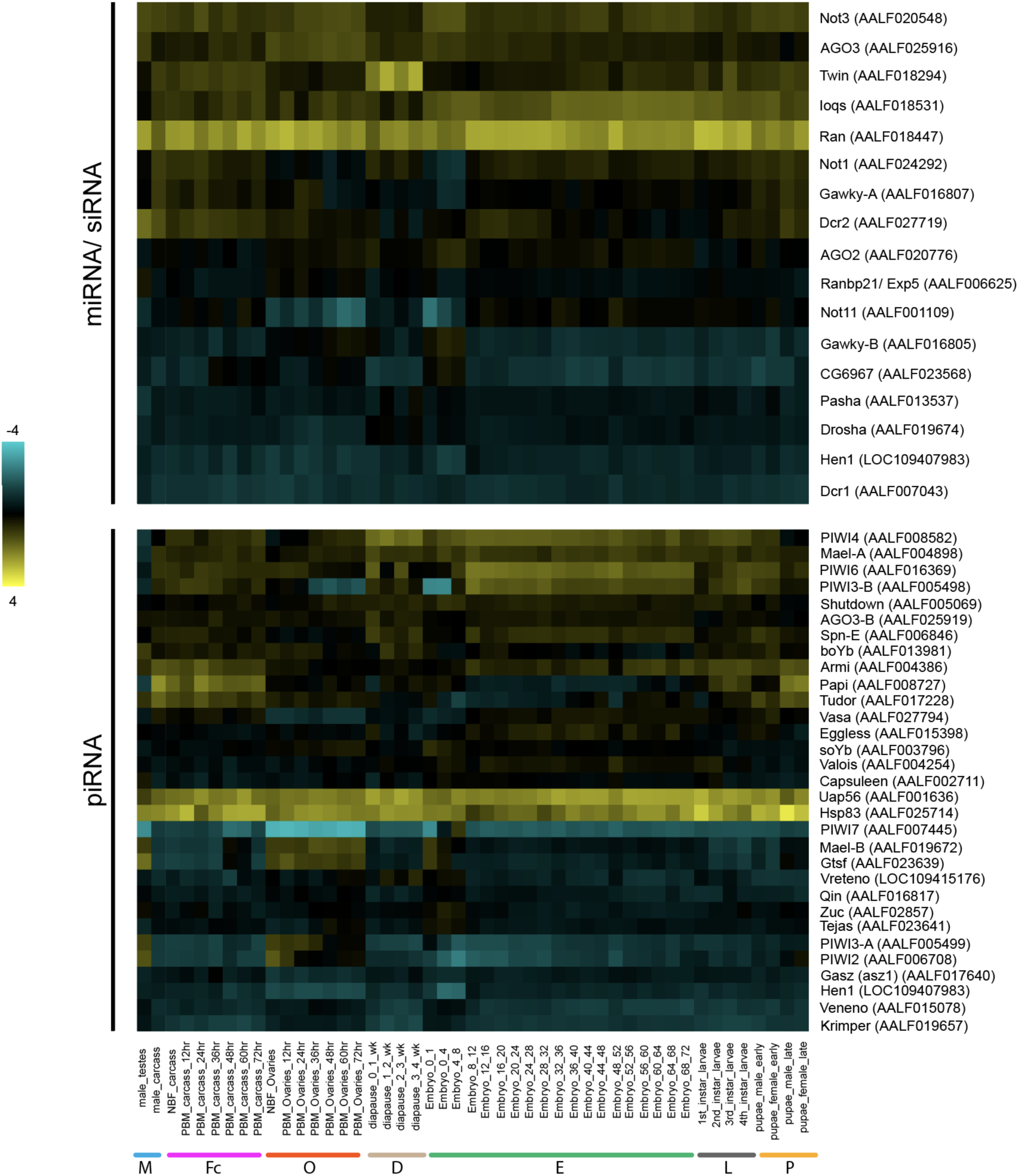
Expression dynamics of protein genes involved in miRNA, siRNA and piRNA pathways. Heat map depicts the z-scores of log2(FPKM + 1) transformed FPKM values. Gene names are listed on the right hand side of the map with their corresponding *Ae. albopictus* AALF IDs in parentheses. Developmental stages are on the bottom of the maps.

**Supplementary Figure 5.**
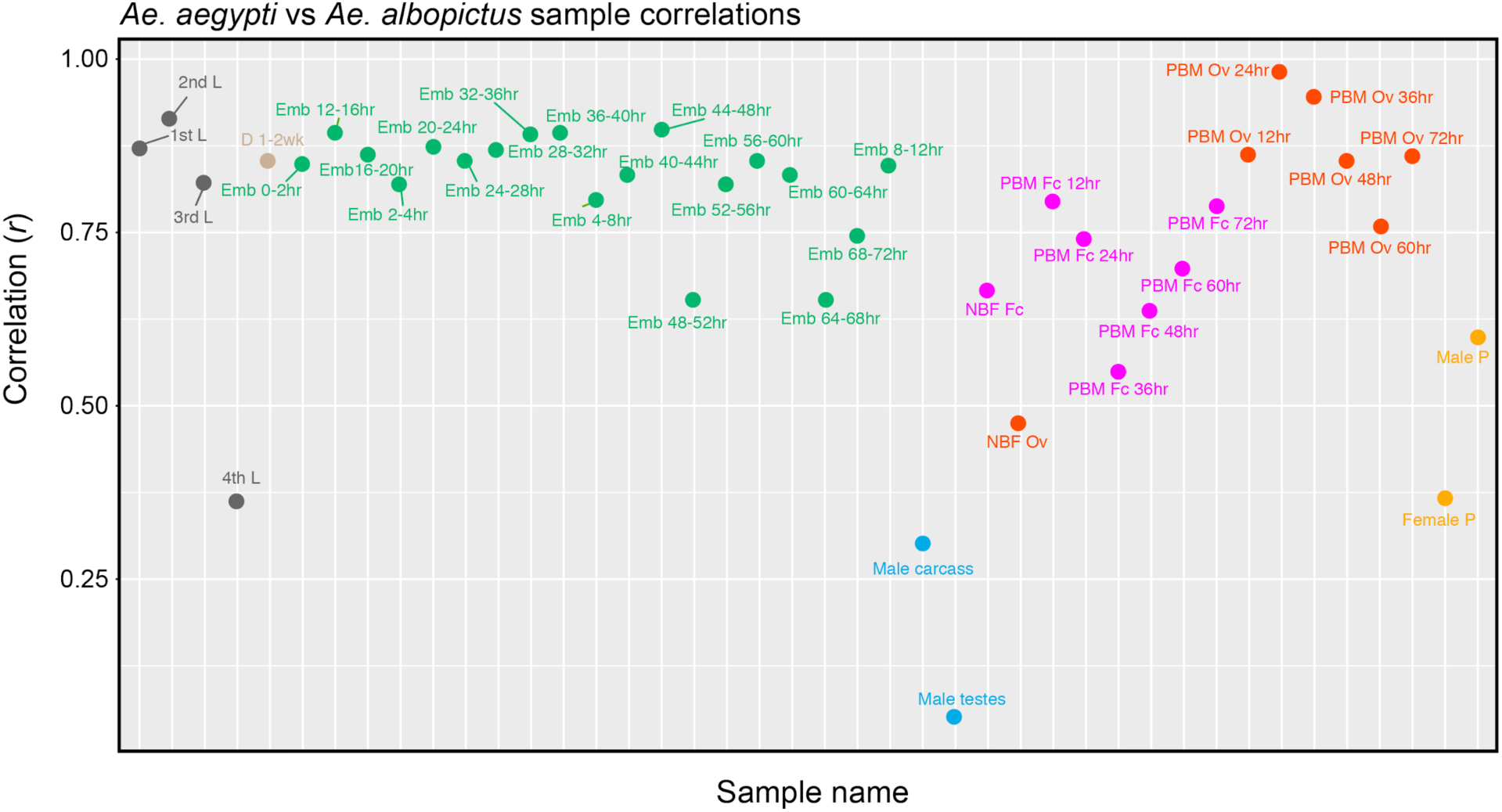
Sample correlations of each developmental time point between *Ae. aegypti* and *Ae. albopictus.* Correlation values were determined by comparing the gene expression of orthologs between the species. The male testes, male carcass, 4th larval instar, and female pupae are highly dissimilar between the two species. The major developmental groups are indicated by color and are organized as follows: M (blue, male testes, male carcass), Fc (pink, NBF carcass, and multiple timepoints PBM: 12hr, 24hr, 36hr, 48hr, 60hr, and 72hr), Ov (orange, NBF ovaries, and multiple ovarian timepoints PBM: 12hr, 24hr, 36hr, 48hr, 60hr, and 72hr), D (tan, diapause at multiple timepoints: 0-1wk, 1-2wk, 2-3wk, and 3-4wk), Emb (embryo at multiple timepoints: 0-1hr, 0-2hr, 2-4hr, 4-8hr, 8-12hr, 12-16hr, 16-20hr, 20hr-24hr, 24-28hr, 28-32hr, 32-36hr, 36-40hr, 40-44hr, 44-48hr, 48-52hr, 52-56hr, 56-60hr, 60-64hr, 64-68hr, and 68-72hr embryos), L (gray, larvae 1st, 2nd, 3rd, and 4th instar larvae stages), and P (yellow, pupae, early male and female, and late male and female pupae stages). Correlation values for each developmental stage are listed in Supplementary Table S16.

**Supplementary Figure 6.**
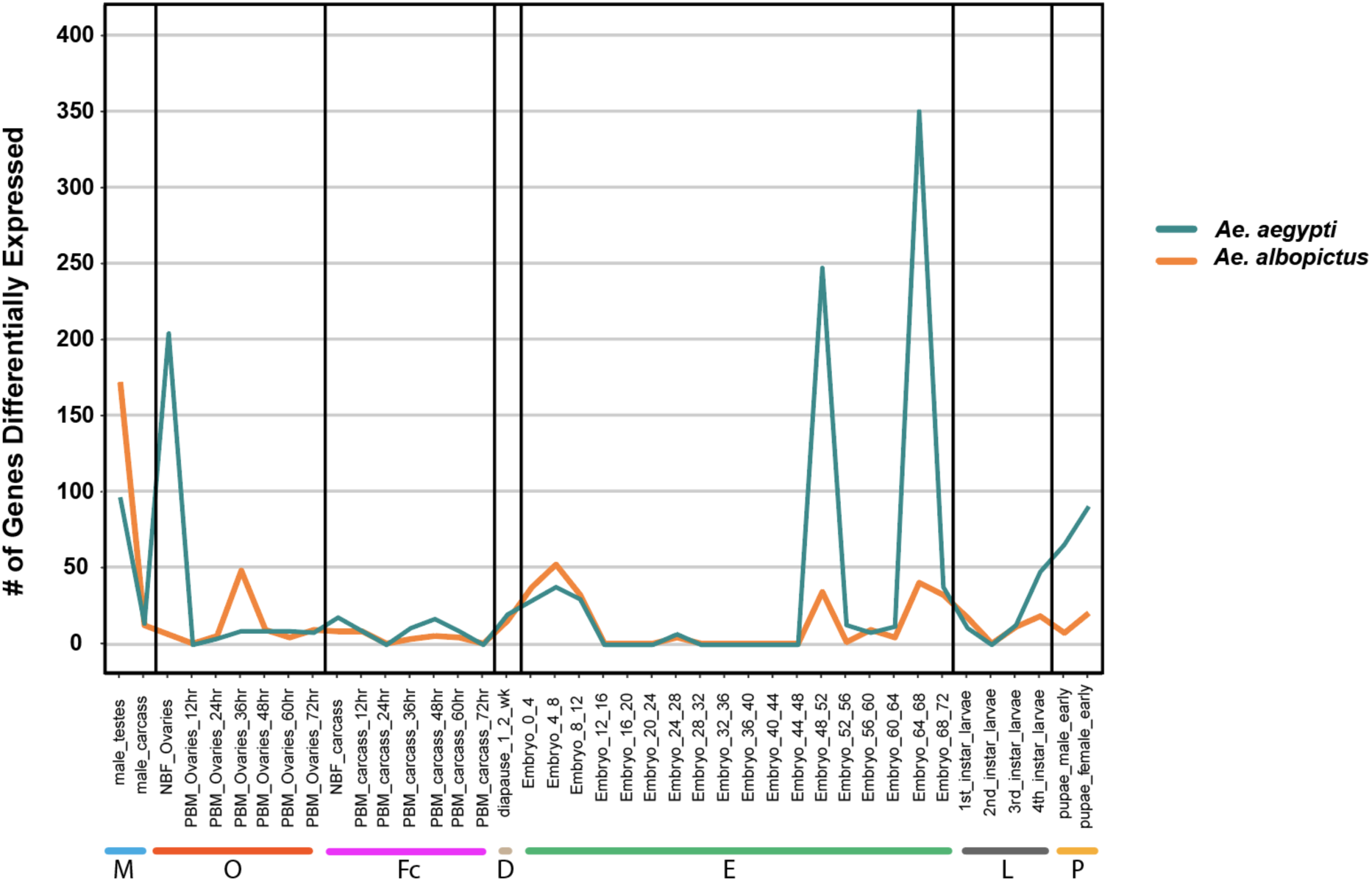
Number of significantly differentially expressed genes on *Ae. aegypti* and *Ae. albopictus* samples across corresponding developmental time points. The number of genes differentially expressed between species were calculated after performing a DESeq analysis on the orthologous genes. Genes with a padj<0.1 and a log2-fold-change of negative (*Ae. aegypti*) or positive (*Ae. albopictus*) values were counted and plotted. The tissues/stages where *Ae. albopictus* differs in gene expression compared to *Ae. aegypti* are the male testes, NBF Ovaries, PBM 36hr Ovaries, late embryonic, and pupal stages. Data can be found in Supplemental Table S17.

**Supplemental figure 7.**
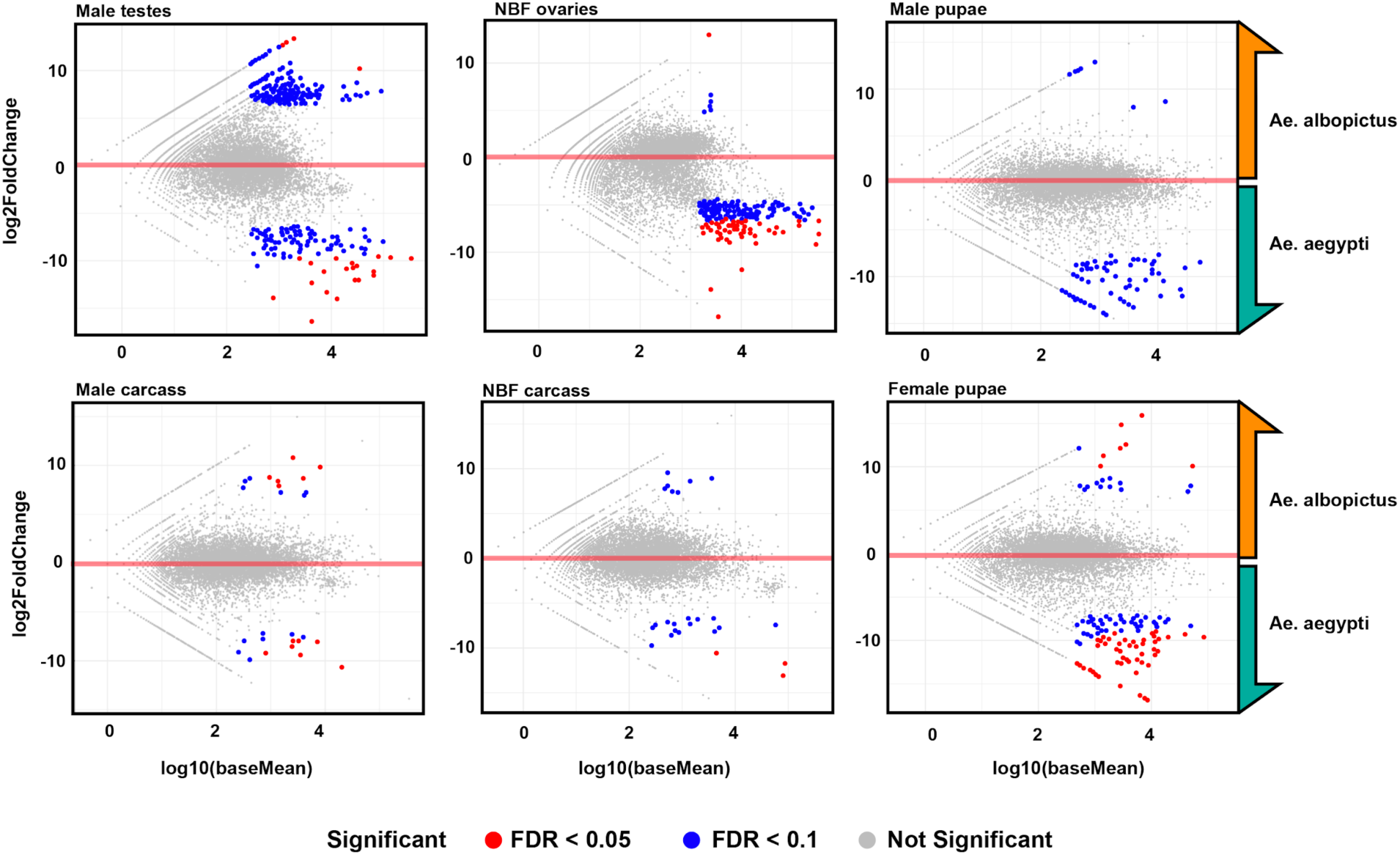
Significant differential expression of orthologous genes in sex specific tissues. Sex-specific tissues analysed were male testes, male carcass, NBF ovaries, NBF female carcass, male and female pupae. The y-axis represents the log2FoldChange and the x-axis represents the log10(baseMean). Genes who show most significant differential expression are colored in red (FDR < 0.05) and genes with significant expression are colored in blue (FDR < 0.1). The red line represents the center of the plot (where log2FoldChange equals 0). The top of the plots indicate genes that are highly expressed in *Ae. albopictus* while genes found below the 0 transparent red line correspond to *Ae. aegypti.* Data can be found in Supplemental Table S17.

**Supplemental figure 8.**
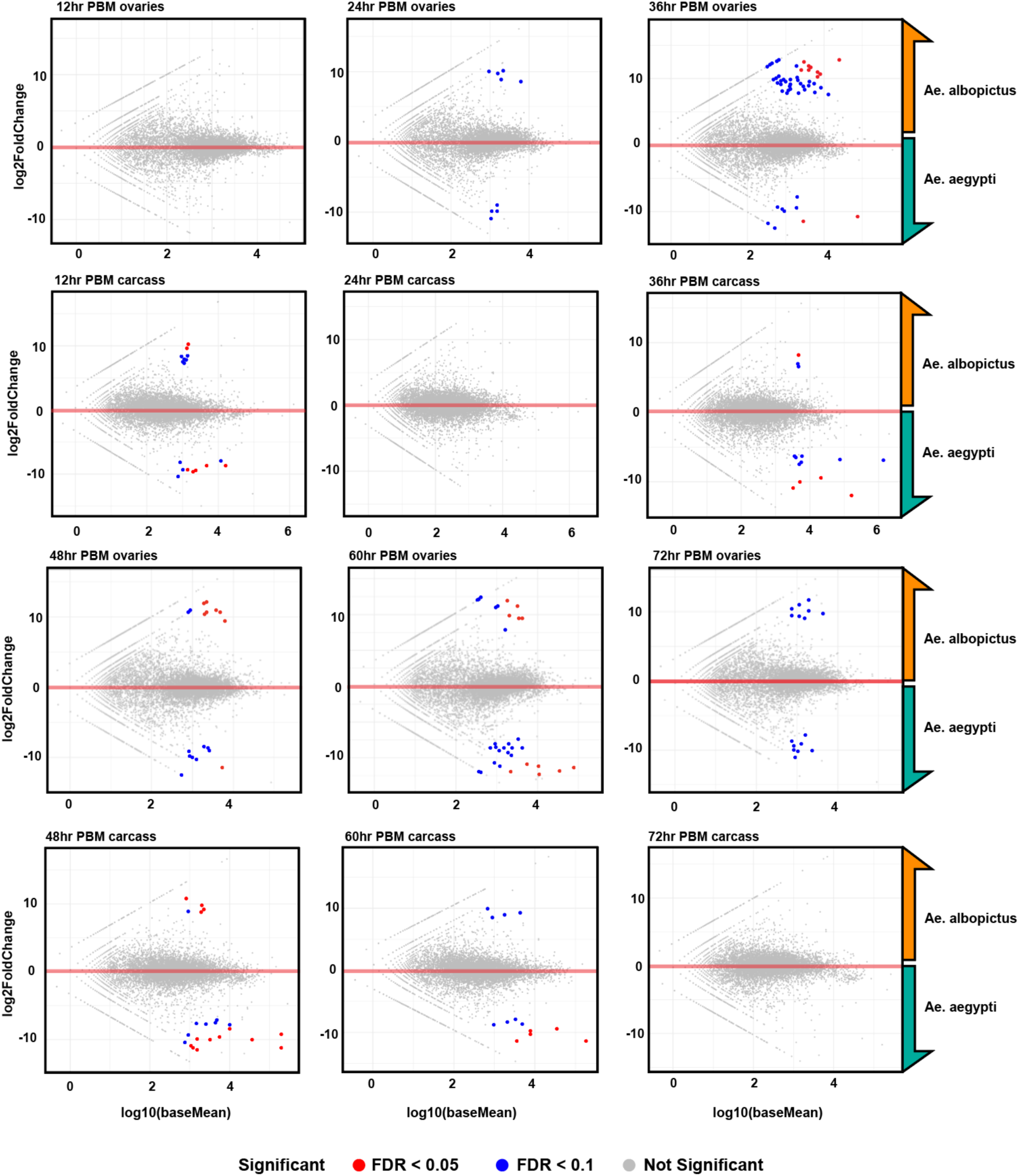
Significant differential expression of orthologous genes in blood-fed female ovaries and carcasses at different 12-hr time points. The y-axis represents the log2FoldChange and the x-axis represents the log10(baseMean). Genes who show most significant differential expression are colored in red (FDR < 0.05) and genes with significant expression are colored in blue (FDR < 0.1). The red line represents the center of the plot (where log2FoldChange equals 0). The top of the plots indicate genes that are highly expressed in *Ae. albopictus* while genes found below the 0 transparent red line correspond to *Ae. aegypti.* Data can be found in Supplemental Table S17.

**Supplemental figure 9.**
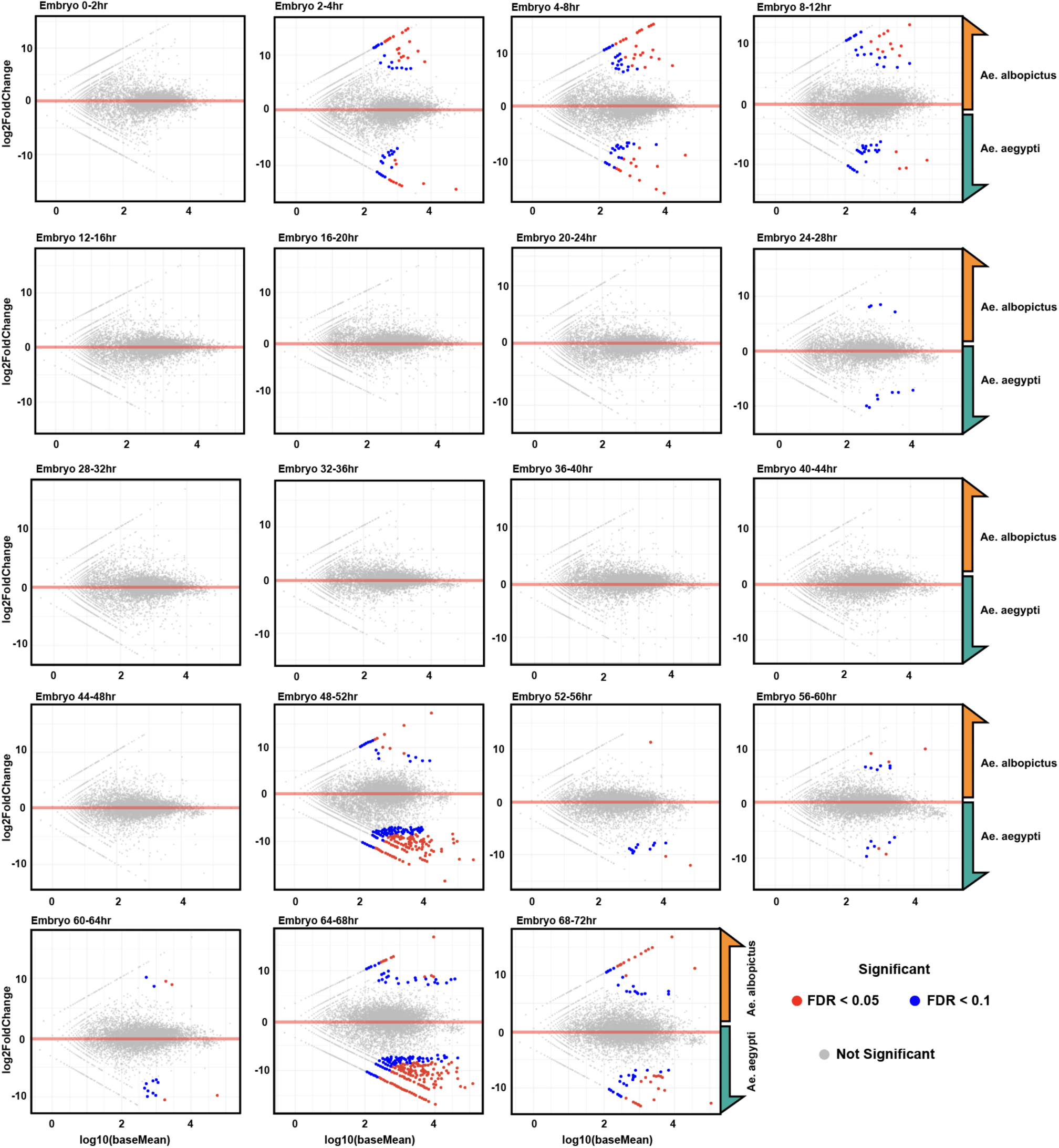
MA plots of differentially expressed genes during embryogenesis of *Ae. albopictus* compared to *Ae. aegypti*. The y-axis represents the log2FoldChange and the x-axis represents the log10(baseMean). Genes who show most significant differential expression are colored in red (FDR < 0.05) and genes with significant expression are colored in blue (FDR < 0.1). The red line represents the center of the plot (where log2FoldChange equals 0). The top of the plots indicate genes that are highly expressed in *Ae. albopictus* while genes found below the 0 transparent red line correspond to *Ae. aegypti.* Data can be found in Supplemental Table S17.

**Supplemental figure 10.**
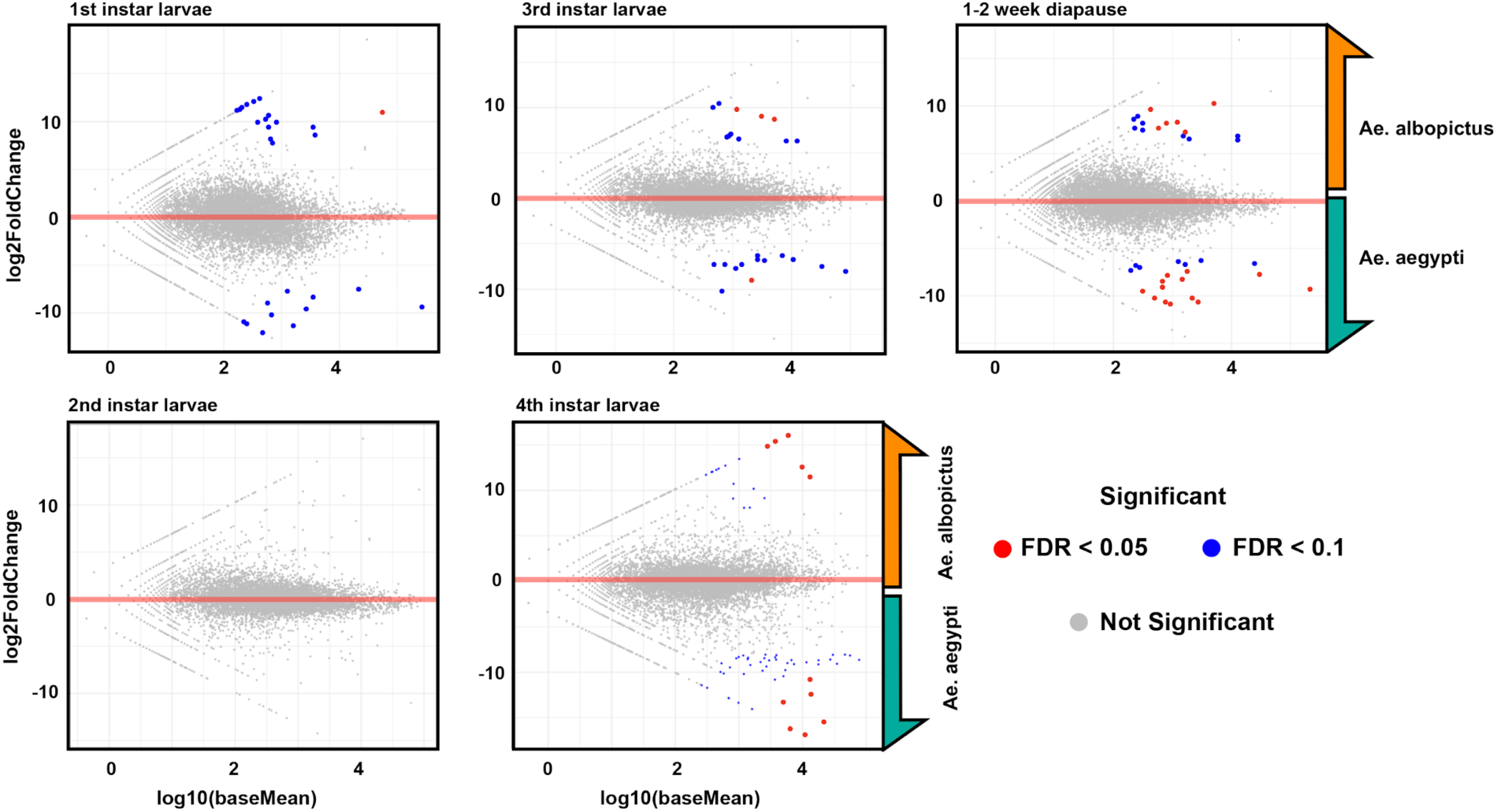
MA plots of differentially expressed genes in the larval instars and diapause samples of *Ae. albopictus* compared to *Ae. aegypti*. The y-axis represents the log2FoldChange and the x-axis represents the log10(baseMean). Genes who show most significant differential expression are colored in red (FDR < 0.05) and genes with significant expression are colored in blue (FDR < 0.1). The red line represents the center of the plot (where log2FoldChange equals 0). The top of the plots indicate genes that are highly expressed in *Ae. albopictus* while genes found below the 0 transparent red line correspond to *Ae. aegypti.* Data can be found in Supplemental Table S17.

